# Theory of temperature-dependent consumer-resource interactions

**DOI:** 10.1101/2020.11.10.376194

**Authors:** Alexis D. Synodinos, Bart Haegeman, Arnaud Sentis, José M. Montoya

## Abstract

Changes in temperature affect consumer-resource interactions, which underpin the functioning of ecosystems. However, existing studies report contrasting predictions regarding the impacts of warming on biological rates and community dynamics. To improve prediction accuracy and comparability, we develop an approach that combines sensitivity analysis and aggregate parameters. The former determines which biological parameters impact the community most strongly. The use of aggregate parameters (i.e., maximal energetic efficiency, *ρ*, and interaction strength, *κ*), that combine multiple biological parameters, increases explanatory power and reduces the complexity of theoretical analyses. We illustrate the approach using empirically-derived thermal dependence curves of biological rates and applying it to consumer-resource biomass ratio and community stability. Based on our analyses, we generate four predictions: 1) resource growth rate regulates biomass distributions at mild temperatures, 2) interaction strength alone determines the thermal boundaries of the community, 3) warming destabilises dynamics at low and mild temperatures only, 4) interactions strength must decrease faster than maximal energetic efficiency for warming to stabilise dynamics. We argue for the potential benefits of directly working with the aggregate parameters to increase the accuracy of predictions on warming impacts on food webs and promote cross-system comparisons.

## Introduction

Temperature strongly regulates consumer-resource interactions that constitute the fundamental blocks of ecosystems (O’Connor *et al.* 2009; Montoya & Raffaelli 2010; Petchey *et al.* 2010; Rall *et al.* 2012; Amarasekare 2019), and anthropogenic climate change will, in most cases, increase mean temperatures (*IPCC 2013*). Therefore, understanding and predicting the impacts of warming on consumer-resource interactions has attracted much interest (Vasseur & McCann 2005; Binzer *et al.* 2012; Thakur *et al.* 2017). A breakthrough occurred with the postulation that metabolic rate increases exponentially with temperature, with the slope (activation energy) conserved across levels of organisation (Gilooly *et al.* 2001; Brown *et al.* 2004). However, activation energies can vary significantly among organisms and biological rates (Dell *et al.* 2011; Réveillon *et al.* 2019). In addition, the thermal response curve of biological rates can decrease at high temperatures, producing a unimodal thermal dependence shape (Deutsch *et al.* 2008; Pörtner & Farrell 2008; Englund *et al.* 2011; Uiterwaal & DeLong 2020). This lack of consensus regarding the exact shape of the temperature-dependence of physiological rates (e.g. ingestion rates), behavioural traits (e.g. consumer search or attack rates) or production (carrying capacity) has contributed to diverging, sometimes contradicting, predictions of how consumer-resource interactions will respond to warming (e.g. Vucic-Pestic *et al.* 2011; Sentis *et al.* 2012).

### A dual approach to address the divergence in predictions

Even though biomass distributions and stability have been much studied properties of consumer-resource communities and food webs (Rall *et al.* 2010, 2012; Uszko *et al.* 2017; Barbier & Loreau 2019; Bideault *et al.* 2020), their predicted responses to warming vary. Biomass ratios have been theorised to increase (Rip & McCann 2011; Gilbert *et al.* 2014) or decrease (Vasseur & McCann 2005) monotonically with warming, though experimentally-derived data have mainly yield unimodal responses (Fussmann *et al.* 2014; Uszko *et al.* 2017). Likewise, the effects of warming on stability remain unclear. Using data on specific rates (e.g. consumer ingestion and metabolism), studies have inferred that stability either increases monotonically (Rall *et al.* 2010, 2012; Vucic-Pestic *et al.* 2011; Fussmann *et al.* 2014) or responds unimodally (Sentis *et al.* 2012; Betini *et al.* 2019) to warming. Theoretical work on stability, in particular the onset of oscillations, expands decades (Rosenzweig & MacArthur 1963; May 1972). Vasseur and McCann (2005) showed that warming will destabilise consumer-resource communities when the consumer metabolic rate increases slower than the ingestion rate. Johnson and Amarasekare (2015) demonstrated the pivotal role of the temperature-dependence of carrying capacity — rather than metabolism and ingestion — in determining warming-stability relationships. All these examples demonstrate that the mixed predictions, whether empirically-derived or theoretical, originate from two distinct sources: the different parameters hypothesised to be driving community responses and the thermal dependence shapes of these parameters. To improve the accuracy of predictions regarding the effects of warming on consumer-resource communities, we need to establish which biological parameters drive community properties (biomass distribution, stability) and to acquire a mechanistic understanding of how their thermal dependence shapes affect community properties.

A dual approach utilising sensitivity analysis and the application of aggregate parameters can address both these issues (Fig. 1). In this study, we illustrate this dual approach using the popular Rosenzweig-MacArthur model (Rosenzweig & MacArthur 1963), although the combined approach of a sensitivity analysis and parameter aggregation is not restricted to this model.

**Figure 1.**
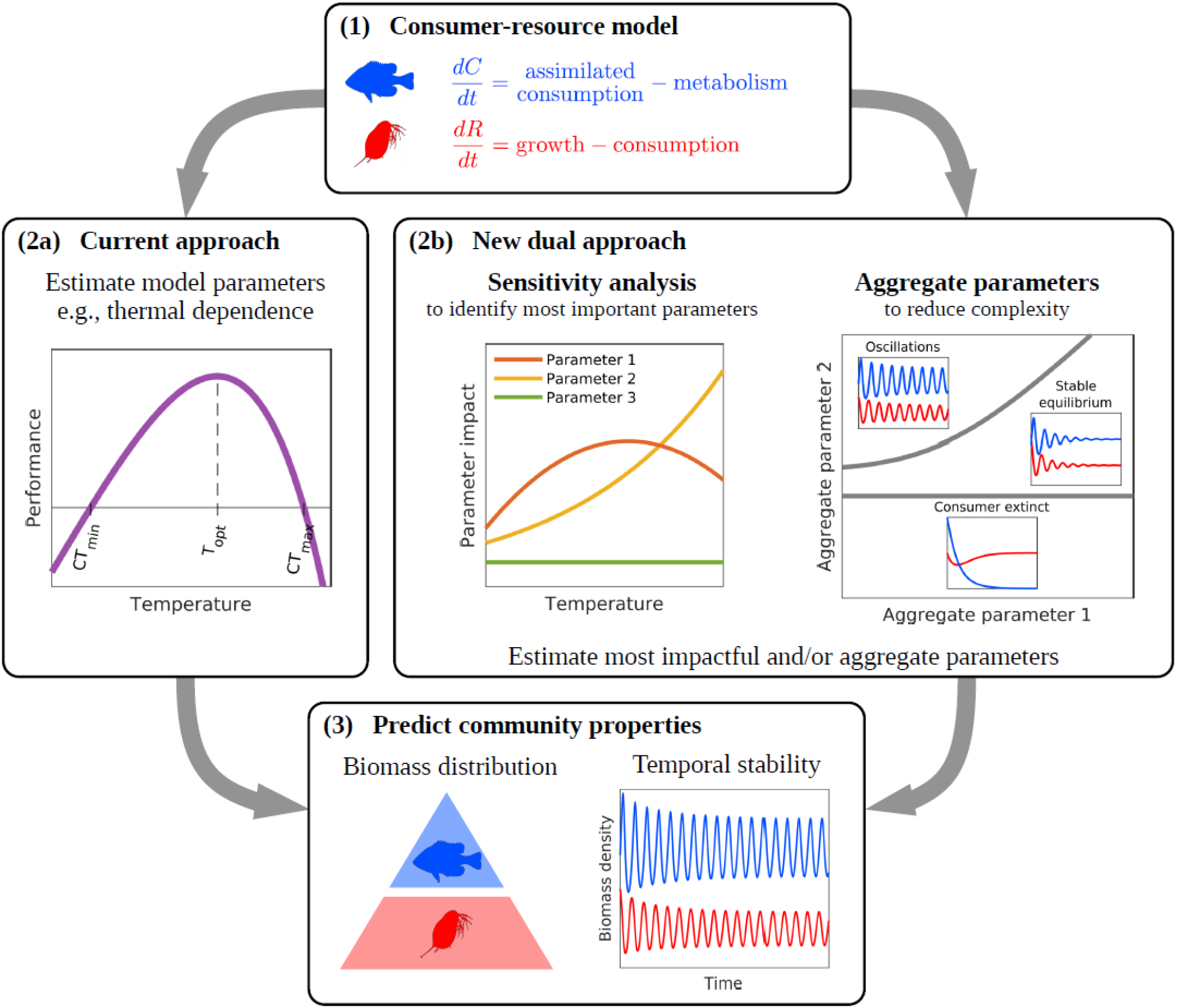
Illustration of the current and new dual approaches to predict the impact of global change drivers on community properties. 1) Predictions require a consumer-resource model; the Rosenzweig-MacArthur model (Rosenzweig & MacArthur 1963) or its bioenergetic equivalent (Yodzis & Innes 1992) have been used most commonly for ectotherm consumer-resource pairs. 2a) The current approach is to experimentally measure the response of parameters along an environmental gradient, e.g. the thermal dependence of the resource population maximal growth rate with critical temperatures, *CT_min_*, *CT_max_*, and the thermal optimum, *T_opt_*. These measurements are used to parameterise the model. Importantly, not all parameters are measured, but rather those which are considered significant (e.g. consumer feeding and metabolic rates for warming-stability relationships). Assuming the remaining parameter values, the model is then used to generate predictions. 2b) Our new dual approach aims to increase the accuracy of predictions and facilitate their comparison. First a sensitivity analysis determines which parameters have the greatest relative impact on the community property of interest along the environmental gradient. Then, aggregate parameters which represent biologically measurable quantities are used to express all sensitivities and determine the dynamics. Collapsing analyses to the two aggregate parameters reduces complexity and increases mechanistic tractability. This facilitates the choice of which parameters need to be measured. 3) Through the empirical determination of the most appropriate parameters (either from the original model parameters of the aggregate parameters themselves) and the reduction in the number of measurements required, prediction accuracy improves. The advantages of the new dual approach are twofold. First, as the sensitivity analysis will have identified the most impactful parameters, the source of divergence in predictions can be isolated. Second, the aggregates represent standardised measurable population-level indicators across systems, making theoretical or empirical predictions directly comparable.

The dual approach benefits from tackling the problem of mixed predictions from different angles. On the one hand, sensitivity analysis establishes the parameters that most strongly influence the community property of interest: it quantifies the increase in a response variable with respect to a small increase in a parameter. It has been extensively used in population ecology and demography (Caswell 2019), with implications for applied ecology (Manlik *et al.* 2018). Since the relative importance of parameters can change along the temperature gradient, a sensitivity analysis allows us to determine the temperatures at which changes in the values of different parameters have the strongest relative impact (Zhao *et al.* 2020).

On the other hand, we can aggregate groups of the primary parameters into fewer, biologically meaningful and empirically measurable quantities. The use of such aggregate parameters reduces the complexity of theoretical analyses, provides a mechanistic interpretation for the difference in predictions and facilitates the comparison among predictions (Barbier & Loreau 2019; Bideault *et al.* 2020). Experimentally, replacing multiple measurements of individual parameters with measurements of the aggregates could also restrict the room for divergent findings. The seminal work of Yodzis and Innes (1992) reduced the analysis of consumer-resource interactions to two aggregate parameters; consumer maximal energetic efficiency and a measure of resource abundance. A variation of maximal energetic efficiency (termed energetic efficiency) has been widely used in empirical studies. However, rather than being measured directly, energetic efficiency has been derived from measurements of its principle components, i.e., feeding and metabolic rates (Rall *et al.* 2010; Vucic-Pestic *et al.* 2011; Sentis *et al.* 2012). Gilbert et al. (2014) posited that interaction strength alone - defined as the impact of the consumer on the resource population density - could capture the effects of warming on the stability of consumer-resource interactions. However, their approach was based on a type I (non-saturating) functional response, whereas most consumer-resource species pairs typically produce type II or III (saturating) functional responses (Jeschke *et al.* 2004). Moreover, the thermal dependence of interaction strength did not match the impact of warming on stability for type II or III functional responses (Uszko *et al.* 2017), pointing to a more complex relationship between interaction strength, warming and stability. We use two aggregate parameters which govern dynamics in the Rosenzweig-MacArthur model: the maximal energetic efficiency of the consumer population, defined as the ratio of energetic gains through ingestion with no resource limitation (i.e., maximal energetic gains) over energetic losses associated to metabolic demand (Yodzis & Innes 1992) and interaction strength, measured as the ratio of resource population density without consumers to resource population density with consumers (Gilbert *et al.* 2014).

Thus, our dual approach identifies the parameters causing the divergence in predictions through the sensitivity analysis and simplifies complex theoretical explorations and empirical measurements through the two aggregate parameters (Fig. 1). The combination of sensitivity analysis and parameter aggregation can be generally applied as it is not tailored to a specific model of consumer-resource interactions; sensitivity analysis (e.g. Chitnis *et al.* 2008) and parameter aggregation (e.g. Barbier & Loreau 2019) have been used independently for different models. Here, we apply this to the Rosenzweig-MacArthur model (Rosenzweig & MacArthur 1963), a model frequently used to study the effects of temperature on consumer-resource interactions (Fussmann *et al.* 2014; Uszko *et al.* 2017; Daugaard *et al.* 2019; Dee *et al.* 2020). We focus on consumer-resource biomass ratio and a stability metric quantifying the proximity to oscillations; these two variables dominating the literature on the effects of warming on consumer-resource communities (Vasseur & McCann 2005; Rall *et al.* 2008; Uszko *et al.* 2017; Betini *et al.* 2019). We implement different thermal parameterisations from the literature to elucidate how the relative importance of different parameters and their varying thermal dependence shapes impact predicted effects of temperature on consumer-resource interactions. Based on our results, we make four predictions that can be theoretically and empirically tested.

### The dual approach

In this study we illustrate the application of the dual approach (i.e. parameter sensitivity and aggregation) using the Rosenzweig-MacArthur model with a type II functional response (Rosenzweig & MacArthur 1963). This model describes the rate of change in resource and consumer biomass densities:

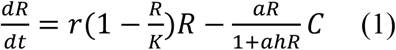

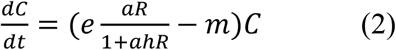

*R* and *C* are the resource and consumer species biomass densities, respectively. Resource growth is logistic, with an intrinsic growth rate, *r*, and carrying capacity, *K*. Resource biomass density is limited by the consumer through a saturating Holling type II functional response with attack rate, *a*, and handling time, *h*. Consumer growth is proportional to the assimilated consumed biomass, with *e* the dimensionless assimilation efficiency; losses occur due to metabolic costs, *m*. Below we present the formulas most relevant to our study; an extensive analysis of the model is available in the supplementary information (SI 1). We chose a type II response due to its prevalence in many natural consumer-resource interactions (Jeschke *et al.* 2004), though our approach works for the general form of the functional response (SI 2). Additionally, the functional response can be defined with respect to attack rate and handing time, 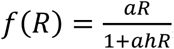, or maximum consumption rate, *J*, and half-saturation density, *R_0_*, 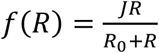 (SI 3).

#### Stability and aggregate parameters

As demonstrated through the ‘paradox of enrichment’ (Rosenzweig 1971), the Rosenzweig-MacArthur model produces population cycles (oscillations) with increasing energy fluxes (Rip & McCann 2011). Therefore, the coexistence equilibrium can be stable or unstable, where dynamics oscillate around the unstable equilibrium (i.e., a limit cycle). The switch from stable to unstable dynamics occurs at a Hopf bifurcation. Theoretical studies have analysed this qualitative change (Yodzis & Innes 1992; Vasseur & McCann 2005; Amarasekare 2015) because these distinct stability regimes translate to different temporal dynamics, with oscillations leading to greater variability over time. We applied a stability metric that quantifies the tendency of the dynamics to oscillate (Johnson & Amarasekare 2015).

For our analyses we assumed dynamics had converged to the stable equilibrium or the limit cycle and determined the coexistence equilibria and the biomass ratio analytically (SI 1). Equilibrium means zero rate of change for both consumer and resource population biomasses. For the limit cycle, this yields the unstable equilibrium which is approximately equal to the time-averaged biomass values along the limit cycle. Thus, we set equations (1) and (2) to zero, solved to yield the coexistence equilibrium and retrieved the biomass ratio:

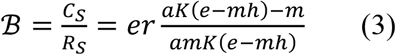

In the model analysis we observed certain repeated parameter groupings (i.e., aggregate parameters) governed the dynamics (SI 1). Such aggregates have been previously used for the analysis of the Rosenzweig-MacArthur model (Yodzis & Innes 1992; Vasseur & McCann 2005). The aggregates we selected represent ecological mechanisms which can be empirically measured. These are maximal energetic efficiency, 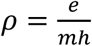, and interaction strength, 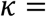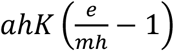. A closer look at *ρ* and *κ* elucidates their biological meaning. 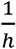 is the saturation value of the functional response and thus represents the maximum consumption rate, *J*. Hence, *ρ* can be written as:

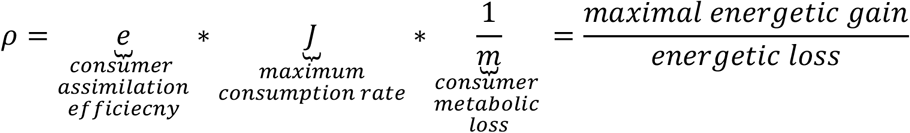

*ρ* quantifies the energetic gain-to-loss ratio of the consumer population biomass assuming its maximum feeding rate is realised (i.e. unlimited resources). *ρ* was introduced by Yodzis and Innes (1992) as a key aggregate parameter to understand food web dynamics. In empirical studies, a variant of *ρ* termed energetic efficiency, *y*, has been often applied (Rall *et al.* 2010; Sentis *et al.* 2012, 2017). Unlike *ρ*, *y* is a function of the full functional response term and hence also depends on resource density, 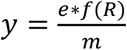, where 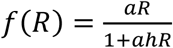 at a specified resource density, *R*.

The second aggregate parameter, *κ*, can be rewritten in terms of the resource population density:

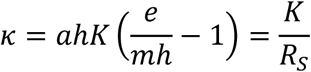

*κ* is the ratio of the resource equilibrium density without consumers (carrying capacity) to the resource equilibrium density with consumers. *κ* quantifies the effect of the consumer population on the resource population and measures interaction strength (Berlow *et al.* 1999; Gilbert *et al.* 2014).

Using *ρ* and *κ* we determine the conditions for positive resource (eq. 4) and consumer (eq. 5) densities, as well as the Hopf bifurcation (eq. 6) (SI 1):

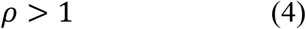

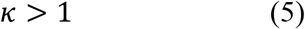

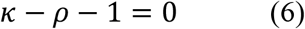

*κ*>1 requires that *ρ*>1 (SI 1). Hence, *κ*>1 defines the consumer feasibility boundary. We do not consider stochastic extinctions which may occur due to large-amplitude oscillations when population biomass reaches very low values.

To determine stability, we adjusted the metric of Johnson and Amarasekare (2015) so that it vanished at the Hopf bifurcation (SI 4). This metric, 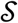, defines stability solely in relation to the Hopf bifurcation.

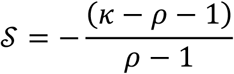

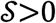 corresponds to a stable equilibrium and 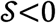 to oscillations.

#### Sensitivity analysis

We performed a sensitivity analysis of the biomass ratio 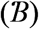 and the stability metric 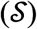 with respect to the original model parameters (i.e., *r*, *a*, *h*, *e* and *m*). A sensitivity analysis quantifies the effect of a small change in a parameter on the response variable. Typically, while one parameter is being perturbed, all others are assumed to remain constant and correlations between the parameters are not explicitly considered. Nevertheless, this approach can be applied in cases of correlated parameter change without a loss of accuracy. When environmental conditions (e.g. temperature) induce correlated changes in the parameters, the sensitivity of the response variable (e.g. 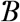 or 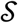) with respect to the environmental conditions can be reconstructed by combining the sensitivities of the individual parameters (SI 5.1). Different types of sensitivity indices exist such as simple sensitivity and elasticity (Manlik *et al.* 2018; Caswell 2019). Here we used elasticity for biomass ratio 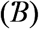 and an adjusted elasticity for stability 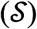 (SI 5.1), both dimensionless to facilitate direct comparisons between parameter sensitivities.

Elasticity is a proportional sensitivity, quantifying how a relative change in a parameter translates into a relative change in the variable; otherwise known as the log-scaled sensitivity (Manlik *et al.* 2018). Thus, the elasticity of 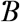 with respect to parameter *x* is given by:

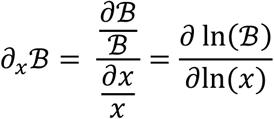

If 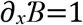, a relative increase of 10% in parameter *x* causes a relative increase of 10% in variable 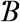. Conversely, 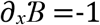 implies that a relative increase of 10% in parameter *x* results in relative decrease of 10% in 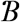.

For the sensitivity of the stability metric, 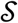, we used a variation of the elasticity. We defined the sensitivity of 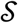 as the incremental change in 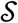 induced by a relative change in parameter *x*. Our adjustment was possible due to 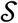 being dimensionless and it prevents sensitivities from diverging to infinity close to the Hopf bifurcation without altering the outcome of our analysis (SI 5.1).

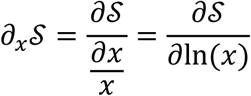

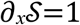 implies that a relative increase in parameter *x* of 10% translates into an absolute increase of 0.1 in 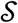 and has a stabilising effect. If, conversely, 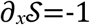, the same relative increase in *x* would lead to decrease of 0.1 in 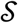 with a destabilising effect.

The magnitude and sign of each sensitivity determine how strongly and in what direction the parameter perturbation impacts the variable, respectively. We used the magnitudes to rank the relative importance of all parameters. The sign provided qualitative information regarding the direction of change (increasing or decreasing the response variable).

All sensitivities could be expressed in terms of *ρ* and *κ*. Hence, sensitivities are fully determined in a plane with of *ρ* and *κ* as axes (Fig. SI 5.1, 5.2). In this *ρ*-*κ* plane the sensitivities of all parameters can be ranked by magnitude, splitting the parameter space into regions where different parameters have the highest, second highest, etc. sensitivity magnitude. Here, we present figures where the regions are determined by the top-two ranked sensitivities; we do not portray changes in the rankings of the lowest sensitivities (see Fig. SI 5.4 and SI 5.5 for the complete biomass ratio and stability metric regions, respectively). The biomass ratio and stability metric have different sensitivity expressions and, therefore, produced different regions. For each variable, the regions remain fixed irrespective of the parameterisation used, because the sensitivity expressions stem from the model equations. The *ρ*-*κ* plane provides additional information, such as the feasibility boundary (eq. 5) and the position of the Hopf bifurcation (eq. 6).

#### Temperature parameterisations

To demonstrate the impacts of different parameter thermal dependencies, we implemented temperature parameterisations from the literature. Maintenance respiration rates, *m*, have been shown to increase exponentially with temperature (Brown *et al.* 2004). The Arrhenius equation is most often used to describe this thermal dependence (Vasseur & McCann 2005; Sentis *et al.* 2017; Uszko *et al.* 2017). However, the thermal response curves of resource growth rate, *r*, attack rate, *a*, and handling time, *h*, have been represented either through the Arrhenius equation (Vasseur & McCann 2005; Binzer *et al.* 2016) or as unimodal functions (Amarasekare 2015; Uszko *et al.* 2017; Zhang *et al.* 2017; Uiterwaal & DeLong 2020; Zhao *et al.* 2020). Carrying capacity, *K*, and consumer assimilation efficiency, *e*, have a less clear connection to temperature (Uszko *et al.* 2017; Dee *et al.* 2020). We selected two parametrisations related to the ongoing debate surrounding the importance of including the decreasing part of the biological rates beyond the optimal temperature (Pawar *et al.* 2016) and used these as an illustrative comparison. The ‘unimodal’ model had a unimodal parameterisation for *r*, *a* (both hump-shaped) and *h* (U-shaped), the Arrhenius equation for *m* (increasing), and constant *K* and *e* (Uszko *et al.* 2017). We compared this to a ‘monotonic’ parameterisation where all thermal dependencies (*r, a, m* increasing; *h, K* decreasing) follow the Arrhenius equation and *e* is constant (Fussmann *et al.* 2014). Following this comparison, we plotted four additional parameterisations from the literature onto the *ρ*-*κ* plane to broaden the comparison and demonstrate the simplicity of applying the approach to empirically-derived measurements. These consisted of two similar monotonic parameterisations (Vucic-Pestic *et al.* 2011; Binzer *et al.* 2016), one where only *a* was hump-shaped (Sentis *et al.* 2012) and one which though monotonic, included some distinctive thermal dependencies – exponentially increasing *K(T)* and *e(T)* and constant *h* (Archer *et al.* 2019). We provide a description of the studies and details of their parameterisations in the supplementary material (SI 6).

We should note that not all parameterisations included the resource growth rate, *r*, so the biomass ratio could not be calculated in these cases. However, we could calculate *ρ* and *κ* and, hence, the biomass ratio elasticities for all parameterisations. Thus, we could determine how biomass ratio sensitivities to individual parameters changed with warming regardless of the actual biomass ratio values. By including studies which had not measured resource growth or estimated the biomass ratio we broadened the scope of the comparison of the biomass ratio sensitivities. Though this does not represent an exhaustive list of parameterisations, we were restricted to parameterisations which could be used to parametrise the Rosenzweig-MacArthur model with a type II response and whose available parameters could yield *ρ* and *κ*. ‘Mild’ and ‘extreme’ temperatures, as well as ‘close’ or ‘far’ from consumer extinction, are defined relative to each parameterisation’s temperature range and feasibility boundaries, respectively. The feasible temperature range was determined by interaction strength (*κ*>1) with the temperature extremes corresponding to the point of consumer extinction. This condition assumes that resources have a broader thermal range than consumers (e.g., Rose & Caron 2007; West & Post 2016). If resources go extinct at temperatures consumers could withstand, then the feasibility boundary becomes dependent on resource growth rate, *r* (Amarasekare 2015). The parameterisations we present come from ectotherms, where environmental temperatures correspond to the organisms’ temperatures. However, our approach can be transferred to endotherms as it does not depend on a specific thermal parametrisation.

### Sensitivities depend on proximity to thermal boundaries

#### Biomass ratio: always most sensitive to e and m

We analytically obtained four groups of biomass ratio elasticity magnitudes, 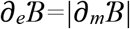,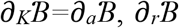 and 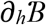. *e* and *m* always have the largest elasticity and hence the strongest relative impact on the biomass ratio (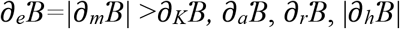, Table 1). Increasing *e* increases the biomass ratio 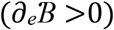, increasing *m* reduces it 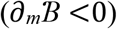. *K* and *a* have equal and positive elasticities 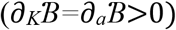, both increasing the biomass ratio. The elasticity of *r* is constant, 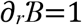; a directly proportional positive effect on 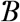. Increasing *h* reduces the biomass ratio, 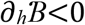. These are general results, independent of any model parameterisation (with temperature or otherwise) following directly from the Rosenzweig-MacArthur model’s equations.

**Table 1.**
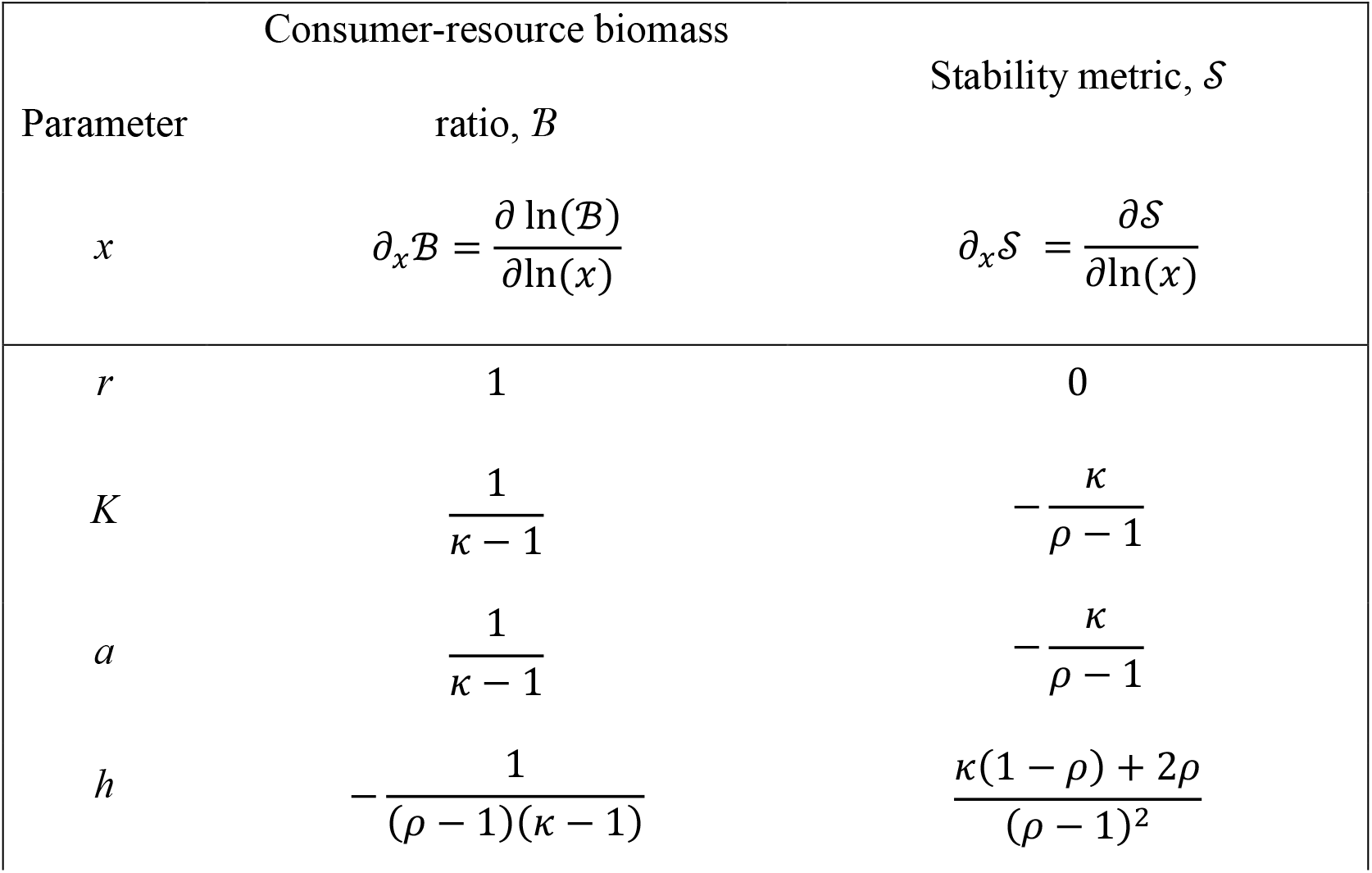
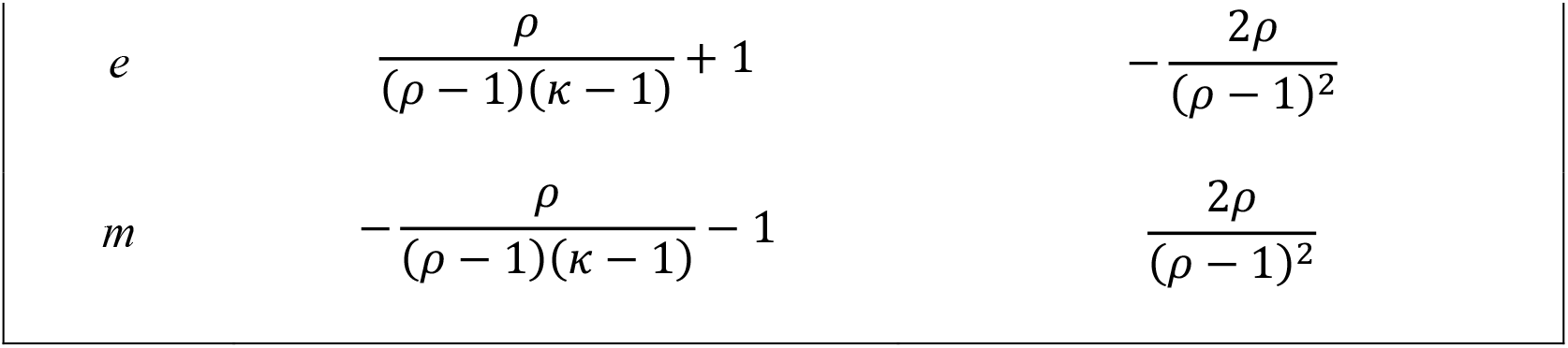
Sensitivities of the biomass ratio 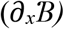 and of the stability metric 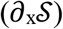 with respect to the six original model parameters. All sensitivities are expressed in terms of *ρ* and *κ*.

#### Biomass ratio: high r elasticity far from thermal boundaries

Both the ‘unimodal’ and ‘monotonic’ temperature parameterisations produced a unimodal biomass ratio thermal dependence (Fig. 2). The unimodal parameterisation induced thermal boundaries to the community at both low (1°C) and high (33°C) temperatures (Fig. 2a). The biomass ratio exceeded 1 for most temperatures (higher consumer than resource biomass), peaked at 14°C around 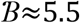 and decreased rapidly to 0 as it approached both thermal boundaries (low and high temperature extremes). The biomass ratio of the monotonic parameterisation (Fig. 2b) increased with warming from low temperatures, peaked at 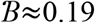, before decreasing to 0 at high temperatures (27.5°C). The two parameterisations are derived from different systems, hence the different temperature ranges.

**Figure 2.**
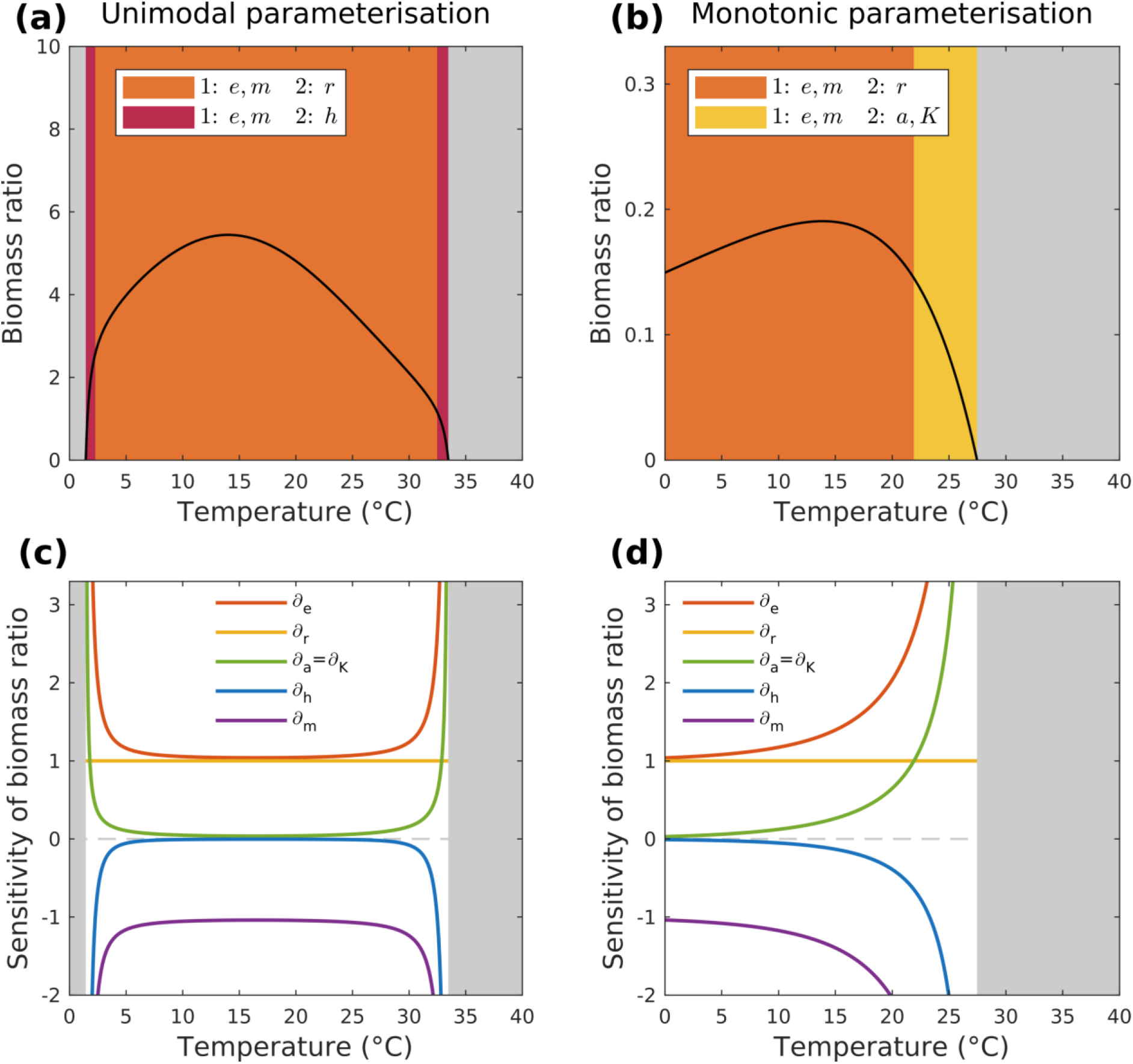
Consumer-resource biomass ratios for the (a) unimodal and (b) monotonic parametrisations along the temperature gradient. Feasible temperature ranges are constrained by the condition of positive biomass densities for both consumer and resource (grey areas correspond to consumer extinction). The different background colours correspond to different elasticity rankings of model parameters (see legend). Panels (c) and (d) provide the values of the six parameter elasticities along the temperature gradient for the unimodal and monotonic parameterisations, respectively.

In both parameterisations, sensitivity to *e* and *m* was highest throughout (Fig. 2c, d) - as expected from the analytical findings. Elasticities were split into two groups at mild temperatures: *e*, *m* and *r* had the highest elasticity with 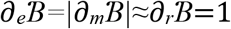, while *a*, *h* and *K* elasticities were very low. Approaching the temperature extremes all elasticities besides 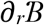 diverged; 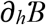 diverged faster than 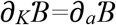 in the unimodal parameterisation (Fig. 2c), while the opposite occurred in the monotonic one (Fig. 2d).

Expressing the elasticities in terms of *ρ* and *κ* (Table 1) reduces the sensitivity analysis to two dimensions. Ranking the elasticity magnitudes creates distinct regions in the *ρ-κ* plane which correspond to different elasticity ranking orderings and provide a mechanistic overview of which elasticities dominate where (Fig. 3a). *e* and *m* always have the highest elasticity, so the three regions reflect changes in the second highest-ranked elasticity. Regions adjacent to consumer extinction (*κ*=1) have high sensitivity to either *h* (red region) or to *a* and *K* (yellow region). *r* elasticity is highest in the region farthest from consumer extinction (orange region). The two temperature parameterisations were mapped onto this plane by calculating their *ρ* and *κ* values (Fig 3b and c). Despite the two trajectories being markedly different, both occupied the region where *r* ranked second highest for most temperatures. The unimodal parameterisation produced a unimodal trajectory, crossing the consumer extinction threshold at low and high temperatures (Fig. 3b). The monotonic parameterisation’s trajectory converged monotonically towards consumer extinction with increasing temperature (Fig. 3c).

**Figure 3.**
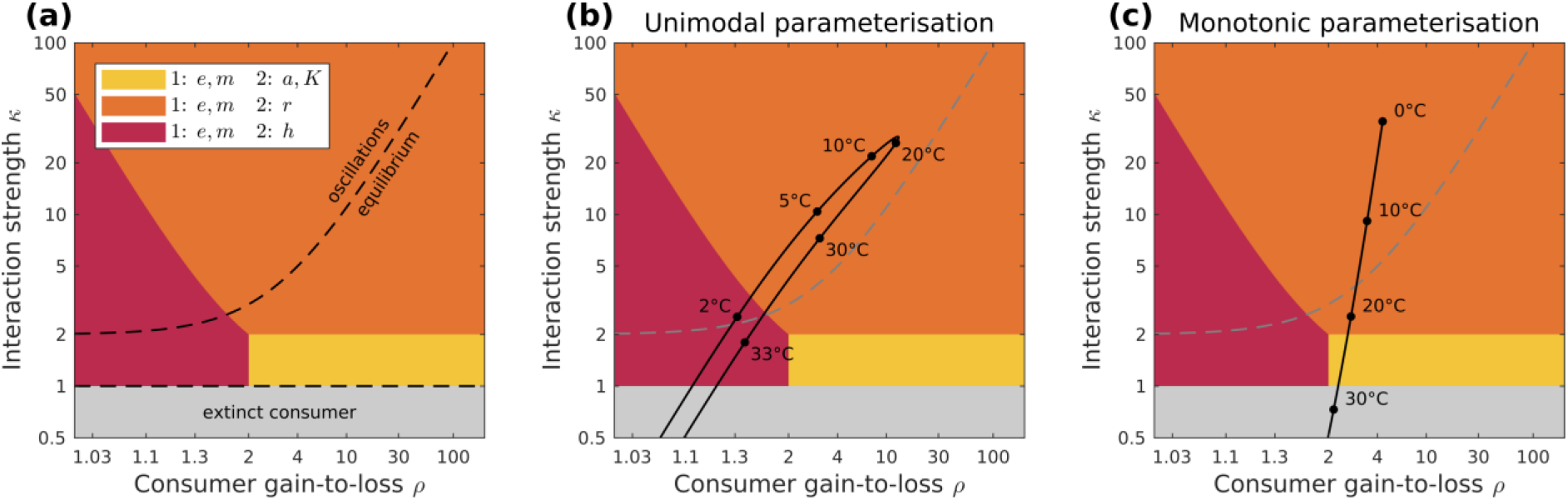
(a) Biomass ratio elasticity rankings in the *ρ-κ* plane. The plane is split into regions (different colours) which correspond to different parameters having the top-two largest elasticities. These regions have been derived from the analytic expressions of the elasticities (Table 1). *e* and *m* elasticities always rank first. Close to consumer extinction *h* ranks second highest at low *ρ* (*ρ* < 2, red region) and *a* and *K* at higher *ρ* (*ρ* > 2, yellow region). *r* ranks second highest far from consumer extinction (orange region). The plane includes the feasibility boundary (*κ*=1) and the Hopf bifurcation (dotted curve splitting the plane into stable equilibrium and oscillations). For the (b) unimodal and (c) monotonic parameterisations from the literature, the thermal dependencies of 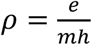 and 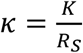 were calculated. This yielded a trajectory for each parameterisation (solid black line). The paths of the trajectories demonstrate the elasticity of the biomass ratio along the temperature gradient for each parameterisation.

All other parameterisations from the literature also occupied the region of high *r* elasticity for most temperatures, far from their thermal boundaries (Fig. 4). Three monotonic parameterisations produced monotonic trajectories (Fig. 4a, b, c) which started in the region of high *r* elasticity and converged monotonically towards consumer extinction (*κ*=1). With a hump-shaped thermal dependence of attack rate, a unimodal trajectory emerged (Fig. 4d). At low temperatures it occupied the region of high *r* elasticity but moved away from consumer extinction. With further warming, the trajectory switched direction and followed the same path as the monotonic parameterisations, crossing the consumer extinction boundary. A unimodal thermal dependence for attack rate (hump-shaped) and handling time (U-shaped) (Fig. 4e), induced extinctions at low and high temperatures imposing a unimodal trajectory. Unlike all previous parameterisations, the trajectory crossed the extinction threshold in the region of high *h* elasticity. The final parameterisation, though monotonic, yielded unimodal trajectories (Fig. 4f). A monotonically increasing *K(T)* (as opposed to decreasing in the other monotonic parameterisations and constant in the unimodal ones) initially forced the trajectory away from consumer extinction, albeit within the region of high *r* elasticity. Consumer energetic efficiency *ρ* decreased, pushing consumers towards extinction, thus forcing an abrupt decline towards the consumer boundary.

**Figure 4.**
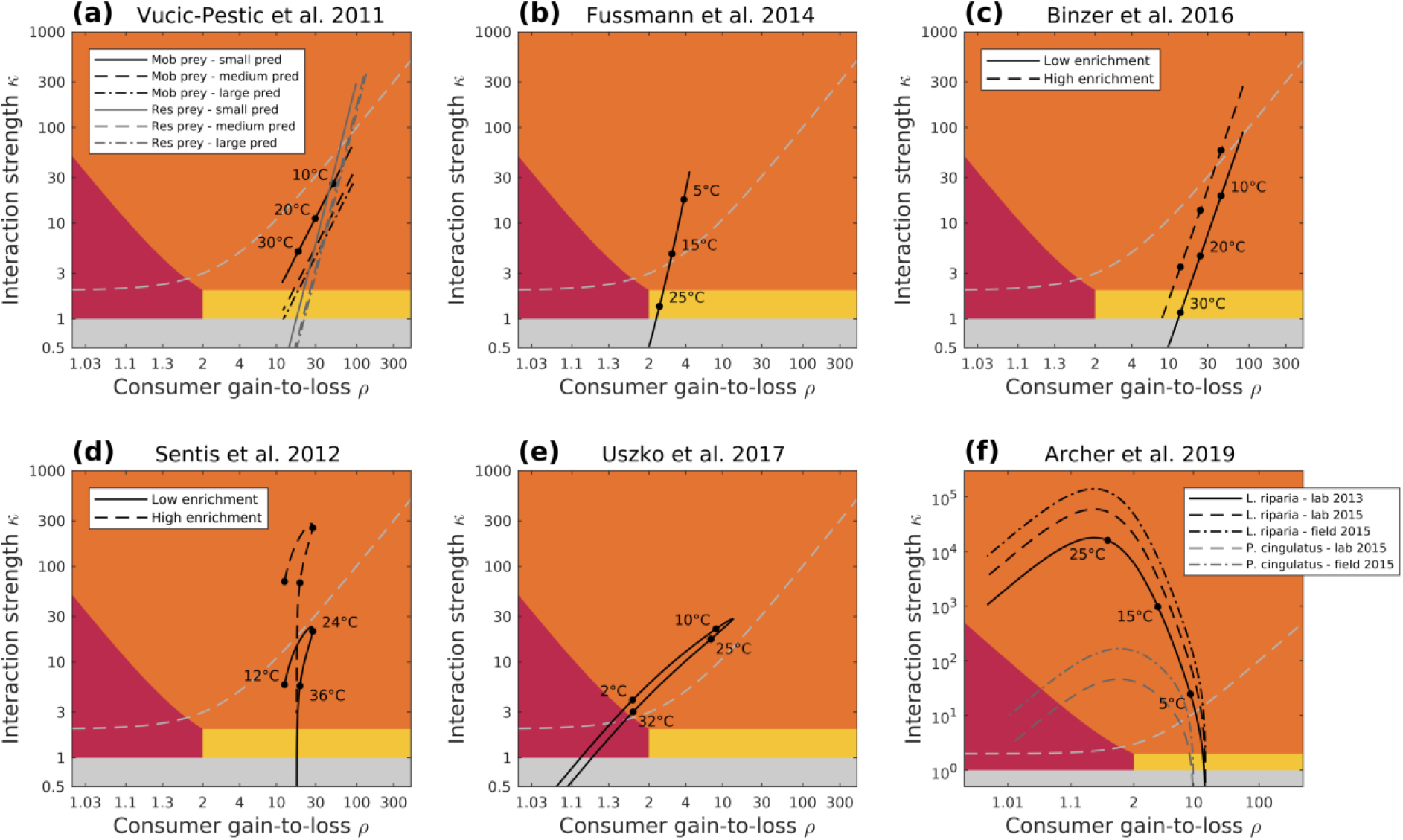
Trajectories of the six empirical temperature parameterisations in the *ρ-κ* plane: a) Vucic-Pestic *et al.* (2011) with six different experiments – three predator size-classes and two types of prey, b) Fussmann *et al.* (2014), c) Binzer *et al.* (2016) with two levels of enrichment, d) Sentis *et al.* (2012), with two levels of enrichment, e) Uszko *et al.* (2017), f) Archer *et al.* (2019) with two prey types and three measurements. (a), (b) and (c) have monotonic thermal dependences for *a, m* (increasing) and *h, K* (decreasing), and a constant *e*. (d) has a unimodal thermal performance curve for *a* (hump-shaped), constant *e* and *K*, monotonic *h* (decreasing) and *m* (increasing). (e) has a unimodal (U-shaped) *h* and *a* (hump-shaped) thermal dependence, constant *e* and monotonic *K, m* (increasing). (f) has monotonic *a, K, m, e* (increasing) and *h* constant. All parameter values are detailed in SI 6. The coloured regions demonstrate the different biomass ratio sensitivity rankings (see legend in Fig. 3a). The trajectories (solid black lines) for each parameterisation are derived from calculating the thermal dependence of 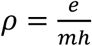 and 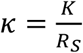 (see *Temperature dependencies and parameterisations* for details).

#### Stability most sensitive either to e and m or to a and K

Similarly to the biomass ratio, the analytical approach for the stability sensitivities yielded results conserved independently of the temperature parameterisations (Table 1): equal sensitivity magnitudes pairwise for *e* and *m* and for *a* and *K* (i.e., 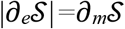 and 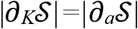), negative stability sensitivities of *e*, *a* and *K* 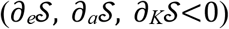 implying they destabilise dynamics, a positive sensitivity of *m* 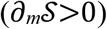 indicating a stabilising effect. *h* can be either stabilising or destabilising and *r* does not affect the stability regime 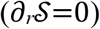.

The unimodal temperature parameterisation exhibited oscillations 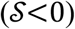 over most temperatures (Fig. 5a). Only at low and high thermal extremes did dynamics briefly stabilise prior to consumer extinction. The monotonic temperature parametrisation produced oscillations at low temperatures 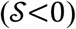, crossed a Hopf bifurcation at 17°C and dynamics were stable 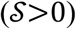 thereafter (Fig. 5b). In both cases, stability close to consumer extinction was most sensitive to consumer assimilation efficiency, *e*, and metabolism, *m*, 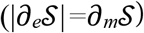 followed by handling time, *h* (Fig. 5c, d). Moving away from the thermal boundaries, attack rate, *a*, and carrying capacity, *K*, increased in relative importance. Furthest away from the thermal boundaries, stability was most sensitive to changes *a* and *K*, followed by *h*. Even though 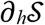 did not rank highest in any temperature range, it was a close second both at the temperature extremes (second to 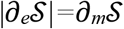) or furthest away from them (second to 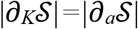). Additionally, *h* switched from destabilising at mild temperatures 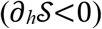 to stabilising 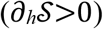 close to the temperature extremes (Fig. 5c, d, Fig. S5.2).

**Figure 5.**
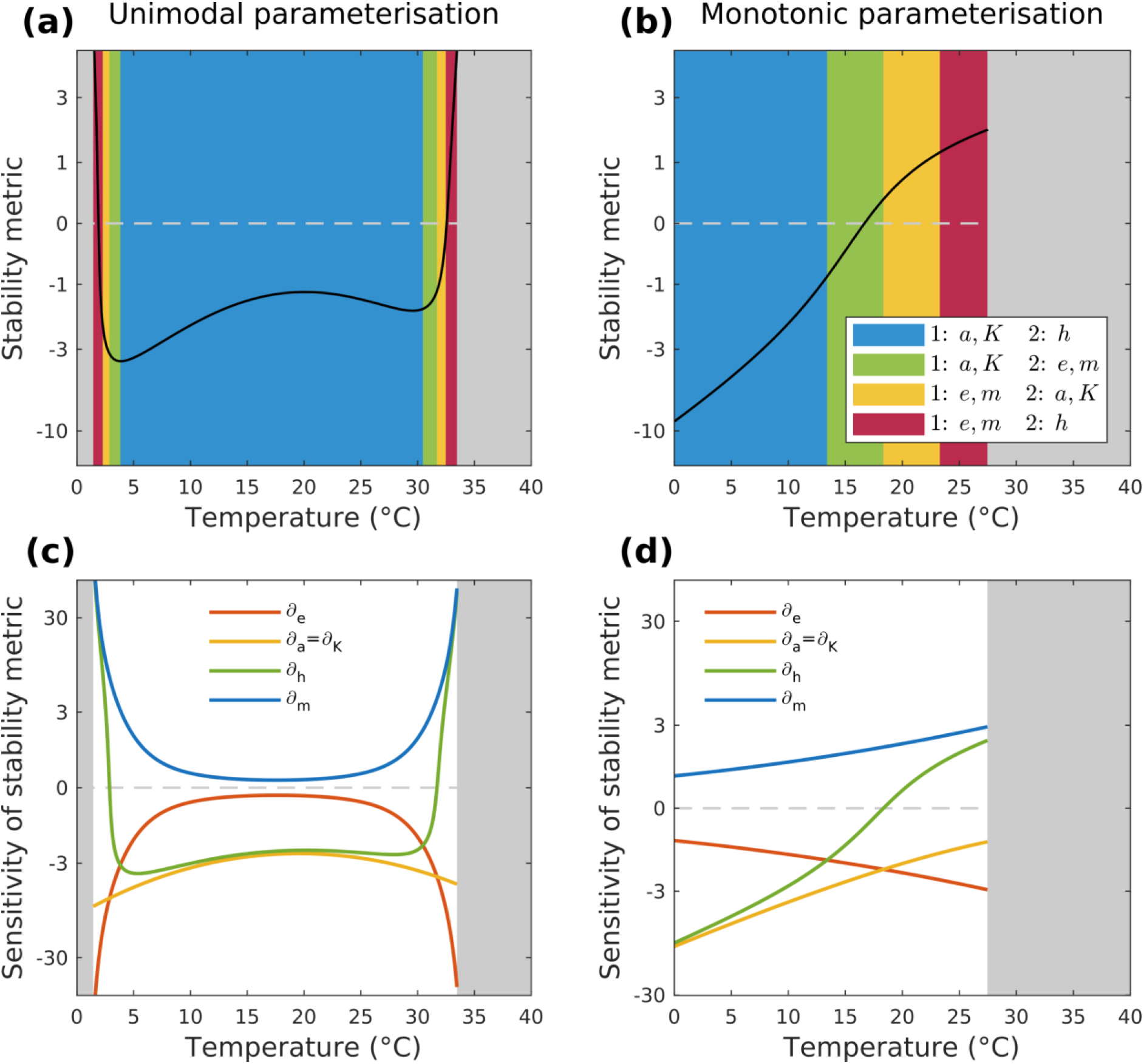
The thermal dependence of the stability metric, 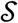, for the (a) unimodal and (b) monotonic parameterisations. 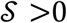 corresponds to stable dynamics, 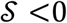 to oscillations. 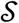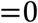 (dotted line) corresponds to the Hopf bifurcation. The coloured temperature ranges highlight regions of different sensitivity rankings. For temperatures beyond the community feasibility boundaries the areas are greyed out. In (c) and (d) the sensitivity to each parameter is plotted along the temperature gradient for the unimodal and monotonic parameterisations, respectively. As resource growth rate does not affect stability (Table 1), we do not plot 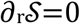.

The *ρ-κ* plane for the stability metric was split into four regions; in two regions closest to consumer extinction, 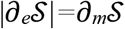 were the largest sensitivities (Fig. 6a, red and yellow regions) and in the two regions furthest from consumer extinction, 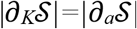 ranked highest (Fig. 6a, green and blue regions). Additionally, the Hopf bifurcation (Fig. 6a, dashed curve) split the plane into stable equilibrium and oscillation regions. Corresponding to the general findings, stability in the two reference (‘unimodal’ and ‘monotonic’) parameterisations was most sensitive to changes in *e* and *m* at the thermal extremes - close to consumer extinction - and to *a* and *K* at milder temperatures – far from consumer extinction (Fig. 6b, c). The unimodal trajectory occupied the region of oscillations for most temperatures, crossing the Hopf bifurcation twice close to consumer extinction, once at low and once at high temperatures (Fig. 6b, blue region). The monotonic trajectory started in the region of oscillations and moved into the stable region with warming, crossing the Hopf bifurcation far from the thermal extreme (Fig. 6c, yellow region).

**Figure 6.**
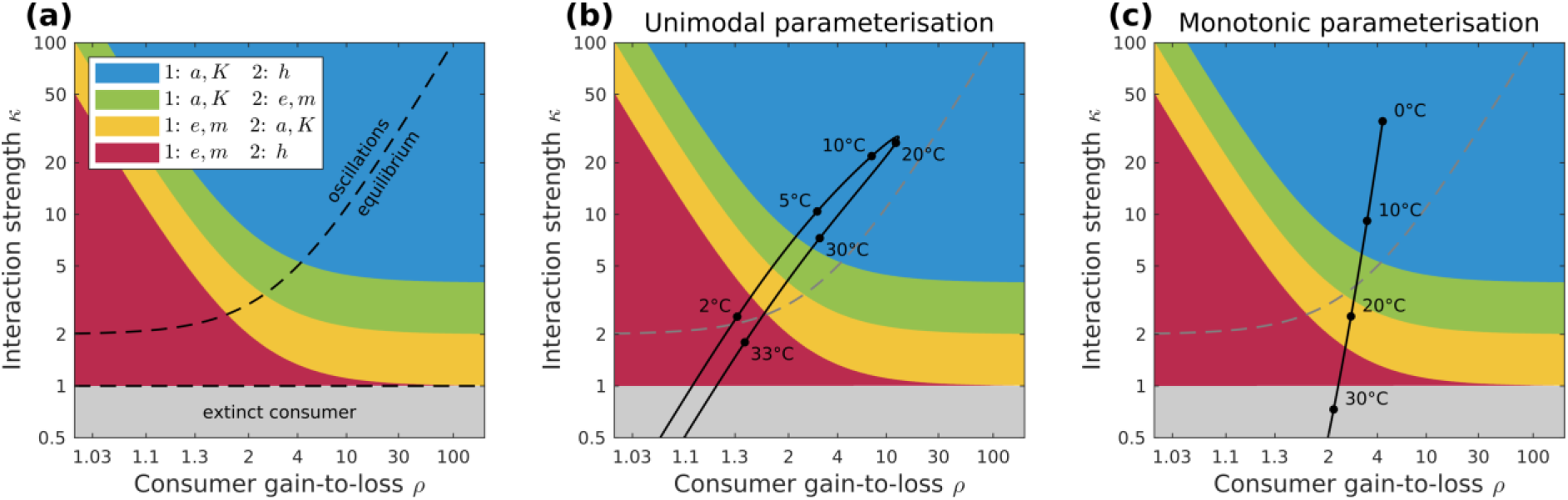
(a) Stability metric sensitivity rankings in the *ρ-κ* plane. The plane is split into regions (different colours) which correspond to different parameters having the top-two largest sensitivities. These regions have been derived from the analytic expressions of the elasticities (Table 1). *e* and *m* rank first close to consumer extinction; *h* ranks second highest closest to consumer extinction (red region). The sensitivity of stability to *a* and *K* increases moving away from consumer extinction. *a* and *K* sensitivity ranks second (yellow region), and then first moving further away. Initially *e* and *m* rank second (green region) before *h* becomes significant (blue region). The plane includes the feasibility boundary (*κ*=1) and the Hopf bifurcation (dotted curve splitting the plane into stable equilibrium and oscillations). For the (b) unimodal and (c) monotonic parameterisations from the literature, the thermal dependencies of 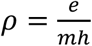 and 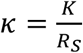 were calculated. This yielded a trajectory for each parameterisation (solid black line). The paths of the trajectories demonstrate the dynamical regime and the sensitivity of the stability metric along the temperature gradient for each parameterisation.

#### Parameter thermal dependencies impact warming-stability relationships

Plotting the other temperature parameterisations’ trajectories onto the *ρ-κ* plane reproduced the same patterns with respect to the stability metric’s sensitivity (Fig. 7): stability was most sensitive to *e* and *m* at the thermal extremes and to *a* and *K* far from the extremes. Significantly, the trajectories revealed the impact of the thermal dependence shape of individual parameters on the warming-stability relationship. In three monotonic parameterisations, warming stabilised the dynamics (Fig. 7 a, b, c). In the cases, when oscillations did take place, these occurred at low temperatures (Fig 7a resident prey, b, c) and dynamics crossed the Hopf bifurcation far from the thermal boundary. In the case with two enrichment levels (Fig. 7c), the high enrichment scenario required higher temperatures to stabilise the dynamics. For the unimodal trajectory with hump-shaped attack rate (Fig. 7d), warming at low temperatures pushed the dynamics towards (low enrichment) or deeper into (high enrichment) the region with oscillations (i.e., destabilised dynamics). Here too, the destabilising impact of enrichment was evident. However, further warming switched the direction of the trajectory. Subsequently, both *ρ* and *κ* decreased. *κ* declined much faster, forcing the dynamics towards the stable region and eventually consumer extinction. Both the switch in the trajectory direction and the Hopf bifurcation (high enrichment scenario) occurred at mild temperatures, in the region of high *a* and *K* sensitivity. In the parameterisation with both *a* (hump-shaped) and *h* (U-shaped) unimodal (Fig. 7e), the Hopf bifurcation occurred close to the thermal boundaries, where *κ* increased (low temperatures) or decreased (high temperatures) much faster than *ρ*. The dynamics were oscillatory for most temperatures, with the switch in the trajectory’s direction occurring in the region of highest sensitivity to *a* and *K*. The final parameterisation’s trajectories were characterised by a negative relationship between *ρ* and *κ* (Fig. 7f). Driven by the positive thermal dependence of carrying capacity, warming increased *κ* and destabilised dynamics which oscillated for most temperatures.

**Figure 7.**
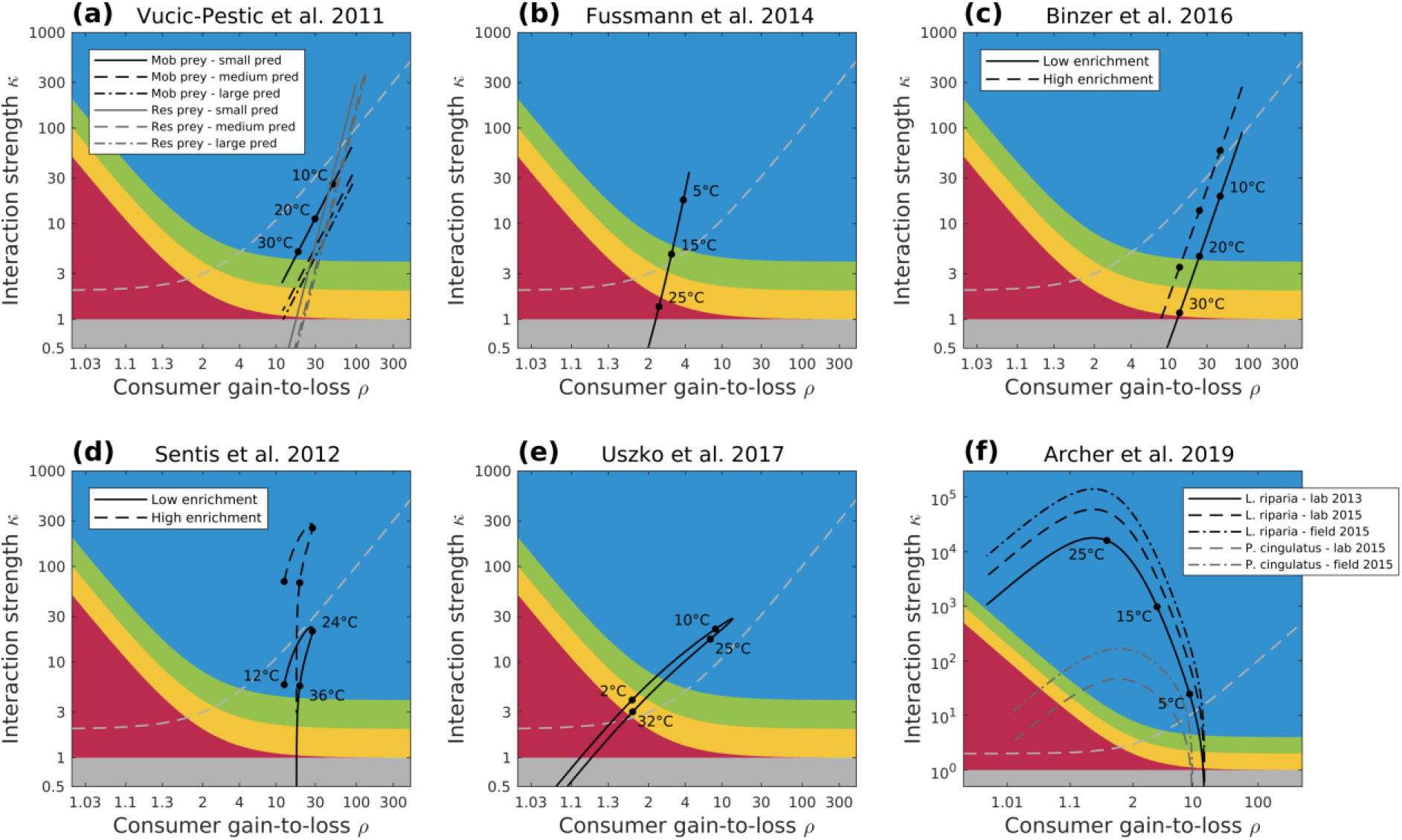
Trajectories of the six empirical temperature parameterisations in the *ρ-κ* plane: a) Vucic-Pestic *et al.* (2011) with six different interaction experiments – three predator size-classes and two types of prey, b) Fussmann *et al.* (2014), c) Binzer *et al.* (2016) with two levels of enrichment, d) Sentis *et al.* (2012), with two levels of enrichment, e) Uszko *et al.* (2017), f) Archer *et al.* (2019) with two prey types and three measurements. (a), (b) and (c) have monotonic thermal dependences for *a, m* (increasing) and *h, K* (decreasing), and a constant *e*. (d) has a unimodal thermal performance curve for *a* (hump-shaped), constant *e* and *K*, monotonic *h* (decreasing) and *m* (increasing). (e) has a unimodal (U-shaped) *h* and *a* (hump-shaped) thermal dependence, constant *e* and monotonic *K, m* (increasing). (f) has monotonic *a, K, m, e* (increasing) and *h* constant. All parameter values are detailed in SI 6. The coloured regions demonstrate the different biomass ratio sensitivity rankings (see legend in Fig. 6a). The trajectories (solid black line) for each parameterisation are derived from calculating the thermal dependence of 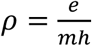 and 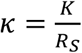. Therefore, trajectories do not change with the variable of interest; the sensitivity regions do.

## Discussion

Research on the impacts of warming on consumer-resource interactions has yielded mixed results (Vasseur & McCann 2005; Englund *et al.* 2011; Rall *et al.* 2012; Gilbert *et al.* 2014; Uszko *et al.* 2017). Resolving this debate and improving predictions has become even more pressing as most ecosystems face increased temperatures (Easterling *et al.* 2000; Walther *et al.* 2002; Root *et al.* 2003; Parmesan 2006). Here, we developed an approach to improve and simplify predictions on the impacts of warming on consumer-resource interactions. This approach integrates two pathways: (1) a sensitivity analysis to identify the key biological parameters whose variations have the largest relative impact on community properties at a given temperature, and (2) aggregate parameters to increase explanatory power. We used the Rosenzweig-MacArthur model with a type II functional response, and applied the approach to consumer-resource biomass ratio and a stability metric quantifying the propensity for oscillations (Johnson & Amarasekare 2015). Therefore, our insights and the aggregates maximal energetic efficiency, *ρ*, and interaction strength, *κ*, apply to study systems well-described by the Rosenzweig-MacArthur model. Our analyses revealed that the relative significance of different parameter groupings is determined by the proximity of the consumer to its thermal boundaries. We, further, elucidated how differences in the shape of the thermal dependence curves of individual parameters qualitatively impact predictions. We used empirically-derived thermal dependence curves of biological parameters from the literature to illustrate this.

We focus our discussion on the formulation of four testable predictions arising from our results. For each prediction, we present its implications and rationale. Then, we discuss the empirical measurement of the aggregate parameters and present important subtleties and potential extensions of our approach.

### Prediction 1: Resource growth rate regulates biomass distribution at mild temperatures

#### Implications

We showed that the relative dominance of consumer assimilation efficiency, metabolism and resource growth rate in driving changes in biomass distributions should manifest itself in any consumer-resource community far from its feasibility boundaries, assuming these communities are well-described by the Rosenzweig-MacArthur model (Fig. 3a). Due to the agreement about the thermal dependence of metabolism (Rall *et al.* 2010; Fussmann *et al.* 2014; Uszko *et al.* 2017) and the negligible -if any- change of assimilation efficiency with warming (Dell *et al.* 2011), differences in the thermal performance curve of resource growth rate will strongly impact biomass ratio predictions. Therefore, improved predictions about the impacts of warming on biomass distributions at mild temperatures necessitate the accurate description of the thermal dependence of resource growth rate.

#### Reasoning

Far from the community thermal boundaries, consumer assimilation efficiency, metabolism and resource growth rate always had the greatest elasticity with an almost equal relative impact on biomass ratio (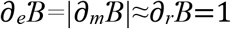, Fig. 2c, d and Fig. S5.1). Increasing metabolism reduced biomass ratios (Table 1), which is likely to be a universal response across ecosystems, given the positive exponential dependence of metabolism on temperature across organisms (Gilooly *et al.* 2001; Brown *et al.* 2004; Rall *et al.* 2012 but see Ehnes *et al.* 2011). Conversely, assimilation efficiency increased biomass ratios but has either been assumed to be unaffected by temperature changes (Vasseur & McCann 2005; Sentis *et al.* 2017; Uszko *et al.* 2017) or has yielded a weak temperature-dependence (Wurtsbaugh & Davis 1977; Handeland *et al.* 2008; Lang *et al.* 2017; Daugaard *et al.* 2019) with negligible change compared to other parameters. Increasing resource growth rate also increased the biomass ratio. However, evidence on the shape of resource growth’s thermal response remains inconclusive: it can either increase exponentially with temperature (Savage *et al.* 2004) or decrease abruptly beyond the thermal optimum (Dannon *et al.* 2010; Thomas *et al.* 2012). Since the biomass ratio is directly proportional to the resource growth rate (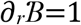, Table 1), it will be strongly affected by the values and shape of the resource growth rate thermal performance curve. Given the consensus surrounding the temperature-dependence of metabolism and the minor scale of potential change in assimilation efficiency with temperature, our findings emphasise the significance of correctly parameterising the resource growth rate when aiming to predict biomass distribution changes due to warming at mild temperatures.

### Prediction 2: Interaction strength determines consumer survival with increasing temperatures

#### Implications

If resources have a broader thermal range compared to consumers (Rose & Caron 2007; West & Post 2016), the thermal boundaries of the community can be determined by measuring solely the thermal dependence of interaction strength, *κ*. This quantity — the ratio of the resource equilibrium density without consumers (carrying capacity) to the resource equilibrium density with consumers — can be determined experimentally (Berlow *et al.* 2004) or through observations, facilitating predictions and cross-system comparisons thereof.

#### Reasoning

Close to the consumer extinction boundary, consumer survival becomes extremely sensitive to all parameters apart from resource growth (Fig. 2, S5.1), making accurate predictions challenging. Currently, consumer survival has been inferred through energetic efficiency — the effective energetic gain of consumers at a certain resource density — which requires determining the thermal dependence of the functional response (Vucic-Pestic *et al.* 2011; Archer *et al.* 2019). Not only is the functional response’s thermal dependence hotly contested (Uszko *et al.* 2017; Uiterwaal & DeLong 2020), but this uncertainty will be exacerbated by its extremely high sensitivity at the community’s thermal boundaries. We showed there exists an alternative, empirically more direct and theoretically more robust metric to determine consumer survival, and hence community feasibility. Interaction strength — the relative values of resource equilibrium without and with consumers (Berlow *et al.* 1999, 2004; Gilbert *et al.* 2014) — provides the necessary condition for consumer survival (*κ*>1), when resources are thermal generalists compared to consumers. This provides an accurate threshold and represents a measurable quantity that can be standardised across experimental designs and study systems (Berlow *et al.* 2004).

### Prediction 3: Warming reduces community stability at low and mild temperatures

#### Implications

This prediction rests on important assumptions: that resources have a broader thermal range, that organisms currently experience temperatures below their optima (Pawar *et al.* 2016) and that the functional response is of type II with a unimodal thermal dependence (Rall *et al.* 2012; Sentis *et al.* 2012; Kuiters 2013; West & Post 2016; Uszko *et al.* 2017; Uiterwaal & DeLong 2020). We deem these assumptions realistic based on the literature; therefore, we argue that consumer-resource interactions at low and mild temperatures will be destabilised by warming. At higher temperatures, warming should always enhance stability.

#### Reasoning

Stability in the context of consumer-resource interactions has predominantly referred to a qualitative distinction between stable and oscillating dynamics (Rosenzweig & MacArthur 1963; Yodzis & Innes 1992; Vasseur & McCann 2005). We based our analysis on an adjusted stability metric which quantifies the tendency of dynamics to oscillate (Johnson & Amarasekare 2015, SI 4). When comparing existing temperature parameterisations, we found that in most monotonic parameterisations (increasing metabolism and attack rate, decreasing handling time and carrying capacity, assimilation efficiency constant), warming always (i.e., monotonically) stabilised dynamics (Fig. 7a, b, c). The single exception arose when warming and carrying capacity increased simultaneously, which destabilised dynamics (Fig. 7f). Carrying capacity has been described as a proxy for enrichment and its destabilising effect has been established whether independently of temperature (Rosenzweig 1971) or as antagonistic to warming (Binzer *et al.* 2016). When at least one parameter in the functional response had a unimodal thermal dependence (i.e., hump-shaped attack rate or U-shaped handling time), this yielded a unimodal warming-stability relationship (Fig. 7d, e). Significantly, the divergence between the unimodal and (most) monotonic parameterisations in the predicted effect of warming on stability manifested itself at low or mild, rather than high temperatures (Fig. 6, 7). This pattern originates in the impact of the parameters with unimodal thermal dependencies on stability. Attack rate is destabilising (Table 1, McCann 2011). Thus, a hump-shaped thermal dependence of attack rate destabilises dynamics with warming below the thermal optimum and stabilises dynamics beyond it. Handling time is stabilising close to the thermal extremes (Fig. 6c, S5.2). A U-shaped handling time will rapidly decrease with warming from low temperatures, which is strongly destabilising; a corresponding steep increase at high temperatures produces a strong stabilising effect. Thus, warming at high temperatures will always be stabilising. However, at lower temperatures, unimodal and monotonic thermal dependencies produce contrasting warming-stability relationships. Therefore, the thermal dependence shape of the functional response combined with the temperatures currently experienced by communities relative to their optimal temperature will determine the impact of warming on stability (Betini *et al.* 2019).

### Prediction 4: Warming stabilises dynamics only when interaction strength decreases faster than maximal energetic efficiency

#### Implications

The combination of *ρ* — the energetic gain-to-loss ratio of consumers given unlimited resources — and *κ* — interaction strength — accurately describes the warming-stability relationship with no recourse to the thermal dependence shapes of individual parameters, the current temperatures relative to the thermal optima, or the proximity to the thermal boundaries of the community. Therefore, differential responses of resources and consumers to warming (Dell *et al.* 2014) will be encompassed by the thermal dependence of the aggregates – assuming the consumer-resource system is well-described by the Rosenzweig-MacArthur model. The Hopf bifurcation condition (eq. 6) dictates that *κ* should decrease faster than *ρ* for warming to stabilise consumer-resource interactions. Thus, measuring *ρ* and *κ* directly can increase the accuracy of warming-stability predictions and simplify cross-system comparisons.

#### Reasoning

Decreasing energetic efficiency or interaction strength have been considered equivalent to increasing stability (Rall *et al.* 2008, 2010; Sentis *et al.* 2012). Thus, estimates of consumer energetic efficiency or interaction strength based on empirically-derived thermal dependence curves of individual rates (e.g. ingestion rate, attack rate, metabolic rate) have been used to infer the impacts of warming on stability (Rall *et al.* 2010, 2012; Vucic-Pestic *et al.* 2011; Fussmann *et al.* 2014). However, this raises two significant issues. On the one hand, even subtle changes in the thermal dependence shapes of individual parameters can yield all possible outcomes (Amarasekare 2015). On the other hand, reducing the analysis of stability to a single aggregate parameter has limitations. Gilbert *et al.* (2014) described the warming-stability relationship with a single aggregate, interaction strength, but their approach was based on a type I functional response and its predictions do not work well in type II or III scenarios (Uszko *et al.* 2017). Johnson and Amarasekare (2015) and Amarasekare (2015) attained a single aggregate parameter to reduce the complexity of their explorations; however, this lacks descriptive power of the dynamics close to the community’s thermal boundaries (SI 4). Our analysis in the *ρ-κ* plane suggests that stability cannot be reduced to a single aggregate parameter nor does a decrease in either one or both of *ρ* and *κ* suffice to stabilise dynamics. In fact, both *ρ* and *κ* can decrease with warming while dynamics become destabilised. A stabilising effect of warming requires not only a concurrent reduction in *ρ* and *κ*, but also the latter to decrease faster. Critically, both *ρ* and *κ* represent biological quantities which can be consistently measured across study systems.

### Working with the aggregate parameters

Working directly with the two aggregate parameters, maximal consumer energetic efficiency, *ρ*, and interaction strength, *κ*, can simplify empirical measurements and improve the accuracy of theoretical predictions, particularly for field data and experiments, as we argue below. To determine the thermal dependence of maximal consumer energetic efficiency and interaction strength, one can measure consumer population growth given unlimited resources and resource population density in presence and absence of consumers at different temperatures, respectively. These measurements can be performed in the lab and the field. Interaction strength is commonly determined in field experiments where consumers are excluded (Berlow *et al.* 2004; Wootton & Emmerson 2005; Novak 2010; Estes *et al.* 2011). Consumer energetic gain-to-loss ratio under effectively unlimited resources is more rarely estimated. However, it can be derived from consumer population net growth and metabolism and mortality, quantities measured commonly in the field and in the lab (Hanson & Peters 1984; Stemberger & Gilbert 1985; Lampert *et al.* 1986). Moreover, confounding factors in field measurements of the population-level aggregates should generate less uncertainty compared to that of measuring multiple individual parameters, where uncertainty propagates and often generates large uncertainty in model predictions (e.g. Sentis *et al.* 2015). Therefore, working directly with the aggregate parameters can be both simpler and lead to more accurate predictions in the field. On the other hand, measuring the individual parameters in the lab has well-established protocols and a history of reliable outputs, with measurements requiring only short-term experiments as opposed to the aggregates.

The choice between measuring the aggregate or the individual parameters will be informed by the questions and objectives of each study. The aggregates describe population-level mechanisms of consumer-resource interactions, while the individual parameters correspond to physiological or behavioural processes of individual organisms scaled up to the population level. As we argued, the aggregates can provide more accurate predictions for field measurements whereas individual parameters can be accurately measured in the lab. This does raise the question whether measurements in a controlled laboratory environment can represent noisier conditions in the field. It would be useful to compare directly measured aggregate parameters to aggregate parameter values derived from the individual parameters to determine how well predictions based on individual rates capture the dynamics of the system. Regardless of the choice, our approach provides the tools for both pathways: studies working with individual parameters will benefit from identifying the most important parameters to measure, while aggregate parameter datapoints can be directly mapped onto the ρ-κ landscape.

### Subtleties and extensions

The sensitivity analysis quantified the sensitivity of the model variables to small parameter changes. Therefore, applying its insights to data should take into consideration the scales of parameters in the temperature range of interest and potential uncertainties in the parameter estimates (Manlik *et al.* 2018). Hence, our argument for the reduced significance of the thermal dependence of assimilation efficiency in driving changes in biomass distributions, despite its high sensitivity.

Regarding the stability of consumer-resource interactions, the *ρ*-*κ* plane helped visualise the stabilising effect of a type III functional response (Fig. S2.1), which has both theoretical and empirical support (Sarnelle & Wilson 2008; Kalinkat *et al.* 2013; Uszko *et al.* 2017; Daugaard *et al.* 2019). For the type II response, the defining role of the functional response (attack rate) and the carrying capacity has been widely documented (Rosenzweig 1971; Amarasekare 2015; Johnson & Amarasekare 2015; Binzer *et al.* 2016); we add the important caveat that this is the case only far from consumer extinction (Fig. 6a).

Finally, relaxing certain assumptions can extend our approach. Considering the scenario where the consumer has a broader thermal niche relative to that of the resource will make the thermal limits of coexistence dependent on resource growth (Amarasekare 2015). Considering the extinction of populations with very low abundances to account for stochastic extinctions can define a realisable coexistence range within the feasible parameter space. Breaking down the original model parameters (e.g. handling time includes the handling and ingestion of prey) could facilitate our understanding of the role of more fundamental physiological processes in the dynamics. Finally, climate change will lead to stronger fluctuations in temperatures (*IPCC 2013*), which have been shown to alter predictions in consumer-resource dynamics (Vasseur *et al.* 2014; Dee *et al.* 2020). This makes the inclusion of temperature variability an important next step.

## Conclusions

Warming will have significant, but as yet uncertain impacts on consumer-resource interactions which underpin the structure and functioning of ecosystems. We presented an approach that will help to improve the accuracy of predictions and reconcile divergent results by facilitating cross-system comparisons. This approach first determines the parameters whose variations have the largest effect on community properties. Second, it simplifies analyses to a two-dimensional plane of mechanistically tractable aggregate parameters. Applying it to the Rosenzweig-MacArthur model, we showed that close to the consumer extinction boundary (i.e., at temperature extremes) both consumer-resource biomass ratio and stability are most sensitive to changes in consumer assimilation efficiency and metabolism. Far from the boundary (i.e., mild temperatures), biomass ratio is most sensitive to resource growth rate, consumer assimilation efficiency and metabolism. This yielded our first prediction, that resource growth rate regulates biomass distributions at mild temperatures. The consensus around the thermal dependence of metabolism and the limited potential impact of warming on assimilation efficiency, underscore the importance of correctly measuring the thermal dependence of resource growth rate. Using the two aggregate parameters (interaction strength and consumer maximal energetic efficiency) also simplified the study of important properties of consumer-resource interactions. From this followed our second prediction, that the thermal boundaries of the community are defined by interaction strength alone. In terms of stability, we demonstrated that a unimodal thermal dependence of attack rate or handling time alters predictions of warming-stability relationships below the thermal optimum, where many organisms may be currently living. Hence our third prediction, that initial increases in mean temperatures will destabilise consumer-resource interactions. Significantly, our approach elucidates how the thermal dependence of stability can be comprehensively characterised by maximal energetic efficiency and interaction strength values. This produced our fourth prediction; a faster reduction of interaction strength than of maximal energetic efficiency with warming is necessary for dynamics to stabilise. Finally, we demonstrated the potential for targeted experiments to measure the thermal dependencies of maximal energetic efficiency and interaction strength to improve predictions. Ultimately, we show that any temperature parameterisation fitted to the Rosenzweig-MacArthur model can be mapped onto the aggregate parameter plane, revealing its stability landscape, providing a mechanistic interpretation for its predictions and allowing for the cross-system comparison of these predictions.

## Acknowledgments

We thank three anonymous reviewers, Ulrich Brose and John Drake for their constructive feedback. This research is supported by the FRAGCLIM Consolidator Grant, funded by the European Research Council under the European Union’s Horizon 2020 research and innovation programme (Grant Agreement Number 726176), and by the “Laboratoires d’Excellences (LABEX)” TULIP (ANR-10-LABX-41).

