## Supplementary Information for "Theory of temperature-dependent consumer-resource interactions"

**Supplementary Information for manuscript entitled:** Theory of temperature-dependent consumer-resource interactions

**Supplementary Information 1: Rosenzweig-MacArthur model with type II functional response analysis**

The Rosenzweig-MacArthur model (Rosenzweig & MacArthur 1963) describes the rate of change in resource and consumer biomass densities:

$$\frac{dR}{dt} = r\left(1 - \frac{R}{K}\right)R - \frac{aR}{1+ahR}C \qquad \text{(S1.1)}$$

$$\frac{dC}{dt} = \left(e\frac{aR}{1+ahR} - m\right)C \qquad \text{(S1.2)}$$

*R* is the resource species biomass density and *C* the consumer species biomass density. Here we present density with respect to volume [$\frac{mass}{volume}$, but densities can also be with respect to area [$\frac{mass}{area}$ which would require an adjustment for the units of carrying capacity and attack rate. Resource growth is logistic, with an intrinsic growth rate *r* [$\frac{1}{time}$, and carrying capacity *K* [$\frac{mass}{volume}$]. Resource biomass density is limited by the consumer through a saturating Holling type II functional response with consumer attack rate $a$ [$\frac{volume}{time*mass}$] and handling time, *h* [$time$]. Consumer growth is proportional to the assimilated consumed biomass, with *e* the dimensionless assimilation efficiency; losses occur due to metabolic costs, *m* [$\frac{1}{time}$.

**Equilibria**:

For the equilibrium we set equations (S1.1) and (S1.2) equal to zero and solve to yield the coexistence equilibrium:

$$R_S = \frac{m}{a(e-mh)} \qquad \text{(S1.3)}$$

$$C_S = er\frac{aK(e-mh)-m}{a^2K(e-mh)^2} \qquad \text{(S1.4)}$$

And biomass ratio:

$$B = \frac{C_S}{R_S} = er\frac{aK(e-mh)-m}{amK(e-mh)} \qquad \text{(S1.5)}$$

Biologically meaningful values require that $R_S$, $C_S>0$ which yielded two conditions for coexistence:

$$e - mh > 0 \qquad \text{(S1.6)}$$

$$aK(e - mh) - m > 0 \qquad \text{(S1.7)}$$

If (S1.6) fails, i.e., $e$-$mh<0$ then (S1.7) cannot be satisfied because $aK(e$-$mh)>m$ will be impossible (all parameters are required to have positive values). Therefore, if (S1.7) holds, it necessarily implies that (S1.6) is also satisfied. Hence (S1.7) is a necessary and sufficient condition for coexistence.

**Stability**:

The Rosenzweig-MacArthur model with a type II functional response exhibits two distinct dynamical regimes: a stable equilibrium and oscillations around an unstable equilibrium. To determine whether the equilibrium (S1. 3, S1. 4) will be stable or unstable, we look at the community or Jacobian matrix which linearises the dynamical system locally.

Let $f_R = \frac{dR}{dt}$ and $f_C = \frac{dC}{dt}$. The the community matrix, *J*, will be defined as:

$$J = \begin{pmatrix} \frac{\partial f_R}{\partial R} & \frac{\partial f_C}{\partial R} \\ \frac{\partial f_R}{\partial C} & \frac{\partial f_C}{\partial C} \end{pmatrix} = \begin{pmatrix} r\left(1-\frac{2R}{K}\right) - \frac{aC}{(1+ahR)^2} & \frac{-aR}{1+ahR} \\ \frac{eaC}{(1+ahR)^2} & \frac{eaR}{1+ahR} - m \end{pmatrix} \qquad \text{(S1. 8)}$$

To determine the stability of the equilibrium, we insert the values of $R=R_S$ and $C=C_S$ into (S1.8):

$$J_S = \begin{pmatrix} \frac{rm}{aeK(e-mh)}(ahK(e-mh) - e - mh) & \frac{-m}{e} \\ \frac{r}{aK}(aK(e-mh) - m) & 0 \end{pmatrix} \qquad \text{(S1.9)}$$

In general, an equilibrium is stable if all the eigenvalues of the Jacobian have negative real part. For a 2D dynamical system this condition is equivalent with the Jacobian having a positive determinant and a negative trace (i.e., the sum of the diagonal elements). For (S1.9) the determinant is always positive, so the equilibrium is stable if and only if the trace is negative. If the trace is positive, *trace*($J_S$)>0, the equilibrium is unstable and the dynamics oscillate around the unstable equilibrium (limit cycles). The point at which the dynamics switch from stable to limit cycles is termed a Hopf bifurcation and this corresponds to solution of *trace*($J_S$)=0:

$$ahK(e-mh) - e - mh = 0 \qquad \text{(S1.10)}$$

The first term in the trace, $\frac{rm}{aeK(e-mh)}$, can be ignored in terms of the sign and thus the dynamical regime. However, this does not mean that it plays no role on the dynamics. In particular, it determines the speed at which the steady state (i.e., the stable equilibrium or the limit cycle) will respond to a perturbation.

**Aggregates**:

We observed that equations S1.6, S1.7 and S1.10 produce certain repeated parameter groupings: $e-mh$ and $ahK$. Yodzis and Innes (1992) and Vasseur and McCann (2005) used them to construct two dimensionless aggregate parameters. Energetic efficiency, $y=\frac{e}{mh}$, and *'inverse enrichment'* $\frac{1}{ahK}=\frac{R_0}{K}$, since $R_0=\frac{1}{ah}$. We used the first one, naming it maximal energetic efficiency, $\rho=\frac{e}{mh}$, and created a synthetic aggregate parameter, $\kappa=ahK\left(\frac{e}{mh}-1\right)$. The latter corresponds to interaction strength, when reformulated to $\kappa=ahK\left(\frac{e}{mh}-1\right)=\frac{aK(e-mh)}{m}=\frac{K}{R_S}$ (Berlow *et al.* 1999; Gilbert *et al.* 2014). All analyses that we performed can be reproduced with $\frac{1}{ahK}=\frac{R_0}{K}$ as the second aggregate parameter. Our choice of $\kappa$ was driven by the stronger connection of this metric to empirical studies, as opposed to $\frac{R_0}{K}$ which is harder to quantify.

Using dimensionless $\rho$ and $\kappa$, we can transform expressions S1.6, S1.7 and S1.10:

$\rho>1$ (S1.11)

$\kappa>1$ (S1.12)

Now, (S1.12) represents the necessary and sufficient condition for coexistence and hence determines the feasibility boundary of the community.

Similarly, S1.10 becomes:

$$\kappa - \rho - 1 = 0 \qquad \text{(S1.13)}$$

**Supplementary Information 2: Framework for general form functional type response**

Here we demonstrate how the framework can be applied to the Rosenzweig-MacArthur model (Rosenzweig & MacArthur 1963) with the general form of the functional response (i.e. not confined to the Holling type II). The model equations are:

$$\frac{dR}{dt} = r\left(1 - \frac{R}{K}\right)R - \frac{aR^q}{1+ahR^q}C \qquad \text{(S2.1)}$$

$$\frac{dC}{dt} = \left(e\frac{aR^q}{1+ahR^q} - m\right)C \qquad \text{(S2.2)}$$

We consider time units in terms of days, mass in terms of grams and space in terms of volume for illustration purposes. Units can be adjusted to the experimental designs as appropriate. $R$ is the resource species biomass density and $C$ the consumer species biomass density $[\frac{g}{m^3}]$. Resource growth is assumed to be logistic, with an intrinsic growth rate $r$ $[\frac{1}{d}]$ and carrying capacity $K$ $[\frac{g}{m^3}]$. Resource biomass density is limited by the consumer through the functional response with consumer attack rate $a$ $[\frac{m^{3q}}{g^q d}]$ and handling time, $h$ $[d]$. Consumer growth is proportional to the assimilated consumed biomass, with $e$ the dimensionless assimilation efficiency, while losses occur due to metabolic costs, $m$ $[\frac{1}{d}]$. $q$ – a dimensionless parameter – determines the shape of the functional response; for the Holling type II response, $q = 1$, for Holling type III, $q = 2$. We should note that the biomass densities and carrying capacity need not be with respect to volume, $[\frac{g}{m^3}]$; the equations apply equally to surface, $[\frac{g}{m^2}]$. This would require the attack rate to have units, $[\frac{m^{2q}}{g^q d}]$.

The coexistence equilibrium for the general form is:

$$R_S = \left(\frac{m}{a(e-mh)}\right)^{1/q} \qquad \text{(S2.3)}$$

$$C_S = \frac{r}{aK}\left(\frac{m}{a(e-mh)}\right)^{1-q}\left(K - \left(\frac{m}{a(e-mh)}\right)^{\frac{1}{q}}\right)\left(\frac{e}{e-mh}\right) \qquad \text{(S2.4)}$$

$$B = \frac{C_S}{R_S} = \frac{r}{aK}\left(\left(\frac{m}{a(e-mh)}\right)^{\frac{1}{q}}\right)^{1-q}\left(K - \left(\frac{m}{a(e-mh)}\right)^{\frac{1}{q}}\right)\left(\frac{e}{e-mh}\right) \qquad \text{(S2.5)}$$

The aggregate parameters are the maximal energetic efficiency, $\rho$, and interaction strength, $\kappa$. $\rho$ is determined as the maximal possible energetic gains of the consumer, which is reflected by the ratio of maximal potential feeding rate to metabolic losses (Yodzis & Innes 1992b). Maximal feeding rate means a saturated functional response (i.e. unlimited resources and no consumer self-limitation). The functional response, $f(R) = \frac{aR^q}{1+ahR^q}$ saturates at $max\big(f(R)\big) = \frac{1}{h}$. Thus, the term for $\rho$ will be:

$$\rho = \underbrace{e}_{\substack{consumer\\ assimilation\\ efficiecny}} * \underbrace{\frac{1}{h}}_{\substack{maximum\\ consumption\ rate}} * \underbrace{\frac{1}{m}}_{\substack{consumer\\ metabolic\\ loss}} = \frac{maximal\ energetic\ gain}{energetic\ loss} \qquad \text{(S2.6)}$$

The interaction strength, $\kappa$, is defined as the ratio of the resource equilibrium in the absence of consumers (i.e. carrying capacity) to the resource equilibrium with consumers (i.e. coexistence resource equilibrium)(Berlow *et al.* 1999; Gilbert *et al.* 2014).

$$\kappa = \frac{K}{\left(\frac{m}{a(e-mh)}\right)^{1/q}} K\left(ah\left(\frac{e}{mh}-1\right)\right)^{1/q} \quad \text{(S2.7)}$$

Biologically meaningful values require that $R_S$ and $C_S$ are positive, which gives us two conditions on the existence of the coexistence steady state:

$$e - mh > 0 \Leftrightarrow \rho > 1 \quad \text{(S2.8)}$$

$$K\big(a(e-mh)\big)^{\frac{1}{q}} - m^{\frac{1}{q}} > 0 \Leftrightarrow \kappa > 1 \quad \text{(S2.9)}$$

As explained in the main text, condition (S2.9) is necessary and sufficient for coexistence. Therefore, the community feasibility boundary is determined by (S2.9).

The Hopf bifurcation condition can be expressed in terms of $\rho$ and $\kappa$ for the general form of the functional response. The Hopf bifurcation occurs when the trace of the characteristic matrix, calculated at the equilibrium values of resource and consumer, becomes zero. The condition is:

$$m^{\frac{1}{q}}\left(K\left(\frac{a(e-mh)}{m}\right)^{\frac{1}{q}} - 1\right)\left(1 - \frac{q(e-mh)}{e}\right) - m^{\frac{1}{q}} = 0 \quad \text{(S2.10)}$$

By introducing $\rho$ and $\kappa$ this becomes:

$$(\kappa - 1)\left(1 - \frac{q(\rho-1)}{\rho}\right) - 1 = 0 \Leftrightarrow \kappa = \frac{\rho(2-q)+q}{\rho(1-q)+q} \quad \text{(S2.11)}$$

Thus, we have shown how the aggregate parameters can be applied to determine the community feasibility threshold (S2.9) and the Hopf bifurcation (S2.11) for the general form of the functional response.

Below we demonstrate that our framework can be applied to the biomass ratio and stability for the general form of the functional response. For the biomass ratio, we derive the elasticities, but for stability no metric exists that covers the general form of the functional response. Therefore, we will illustrate how the qualitative aspects of the framework's approach can still be applied to investigate stability properties in the $\rho - \kappa$ plane.

*Biomass ratio:*

The biomass ratio was determined to be $B = \frac{r}{aK}\left(\left(\frac{m}{a(e-mh)}\right)^{\frac{1}{q}}\right)^{1-q}\left(K - \left(\frac{m}{a(e-mh)}\right)^{\frac{1}{q}}\right)\left(\frac{e}{e-mh}\right)$.

The elasticities with respect to each parameter, $x$, are defined as $\partial_x B = \frac{\partial ln(B)}{\partial ln(x)}$ . Then, the elasticities for the general form functional response will be:

Table S2: Elasticities of the biomass ratio ($\partial_x B$ with respect to the six original model parameters. All sensitivities are expressed in terms of $\rho$ and $\kappa$.

| Parameter | Consumer-resource biomass ratio, $B$ |
|---|---|
| $x$ | $\partial_x B = \frac{\partial ln(B)}{\partial ln(x)}$ |
| $r$ | 1 |

| | |
|---|---|
| $K$ | $\frac{1}{\kappa - 1}$ |
| $a$ | $\frac{1}{q}\frac{1}{\kappa - 1}$ |
| $h$ | $\frac{-1}{q}\frac{1}{(\rho - 1)(\kappa - 1)}$ |
| $e$ | $\frac{1}{q}\frac{\rho}{(\rho - 1)(\kappa - 1)} + 1$ |
| $m$ | $\frac{-1}{q}\frac{\rho}{(\rho - 1)(\kappa - 1)} - 1$ |

As we argued, all elasticities are expressible in terms of $\rho$ and $\kappa$. Interestingly, the important results of our sensitivity analysis hold. In particular, far from the consumer extinction, when $\kappa$ is large, $\partial_K B$, $\partial_a B$,$\partial_h B$ all tend towards 0, $\partial_e B$ and $\partial_m B$ towards 1 and $-1$, respectively. Therefore, $\partial_e B = -\partial_m B \approx \partial_r B = 1$ will be the group of largest elasticities and $e$, $m$ and $r$ will be have the highest relative impact on biomass distributions away from the community boundaries. Similarly, close to the community feasibility boundary, $\kappa = 1$, all elasticities $\partial_e B, \partial_m B, \partial_K B$, $\partial_a B$,$\partial_h$diverge apart from $\partial_r B = 1$ which remains constant.

*Stability*

We have already shown that the condition for the Hopf bifurcation can be expressed solely in terms of $\rho$ and $\kappa$ (S2.11, Fig. S2). The stability metric used in the main study (Johnson & Amarasekare 2015), effectively measures the distance from the Hopf bifurcation in the $\rho - \kappa$ plane. Therefore, any distance metric with respect to the Hopf bifurcation and its sensitivities will be expressible in terms of $\rho$ and $\kappa$, as will its sensitivities.

We see that with increasing $q$, the Hopf bifurcation condition becomes steeper. This means that the area below the curve which corresponds to stable equilibrium dynamics grows and hence that a type III functional response leads to more stable dynamics, as has been previously demonstrated (Uszko *et al.* 2017; Daugaard *et al.* 2019). In terms of the effects of warming on stability, two distinct regions emerge: at low maximal energetic efficiency, $\rho < 2$, changes in interaction strength, $\kappa$, determine whether warming will stabilise (decreasing $\kappa$ or destabilise (increasing $\kappa$) dynamics. At higher values of $\rho$, $\rho > 2$, dynamics are effectively stable.

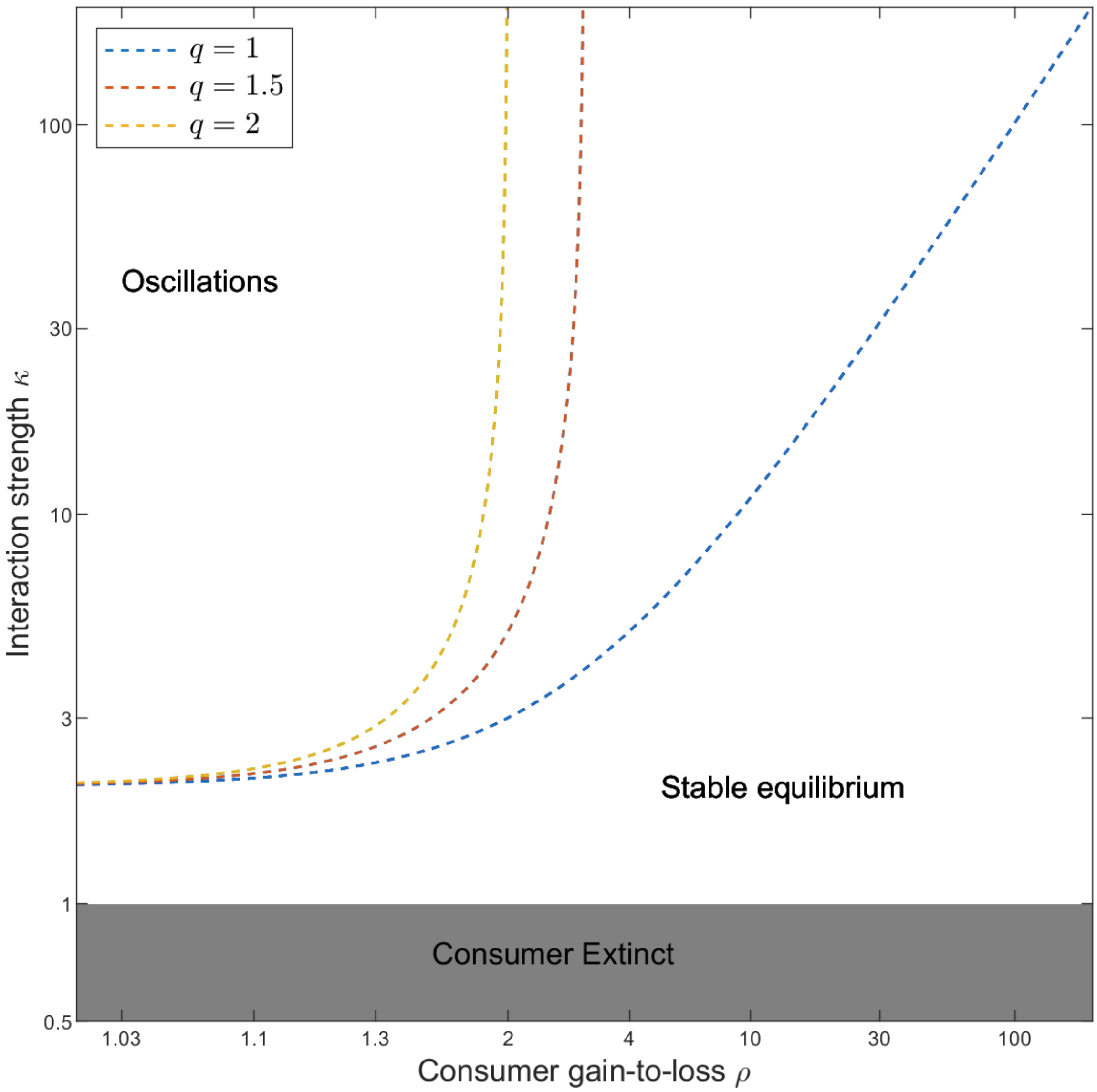

Figure S2 The Hopf bifurcation in the of $\rho - \kappa$ plane for different values of of $q$, i.e. functional response types. $q = 1$ corresponds to the Holling type II and $q = 2$ to the Holling type III. As q increases, the parameter space with stable dynamics expands.

**Supplementary Information 3: type II functional response alternative formulation**

The relationship between consumer per capita feeding rate and resource density, $f(R)$ $[\frac{1}{time}]$, simply known as the functional response can be represented in terms of attack (or search) rate, $a$ $[\frac{volume}{mass*time}]$, and handling time (or inverse maximal intake rate), $h$ $[time]$ (Holling 1959), for a type II response:

$$f(R) = \frac{aR}{1+ahR} \qquad \text{(S3. 1)}$$

Alternatively, the functional response can be reformulated as the Michaelis-Menten equation where the shape of the curve is controlled by the  maximum consumption rate, $J$ $[\frac{1}{time}]$, and half-saturation density, $R_0 \left[\frac{mass}{volume}\right]$ - the resource density corresponding to the point of half-saturation of the functional response, $f(R|0) = \frac{1}{2}max f(R)$. Thus, the functional response becomes:

$$f(R) = \frac{JR}{R_0+R} \qquad \text{(S3.2)}$$

The two formulations are interchangeable through a simple transformation:

$$J = \frac{1}{h} \qquad \text{(S3.3)}$$

$$R_0 = \frac{1}{ah} \qquad \text{(S3.4)}$$

The coexistence steady state becomes:

$$R_S = \frac{mR_0}{eJ-m} \tag{S3.5}$$

$$C_S = erR_0\frac{K(eJ-m)-mR_0}{K(eJ-m)^2} \tag{S3.6}$$

The aggregate parameters:

$$\rho = \frac{eJ}{m}, \kappa = \frac{K(eJ-m)}{mR_0} = \frac{K}{R_S} \tag{S3.7}$$

| Parameter | C:R biomass ratio, $B$ | Stability metric, $\mathcal{S}$ |
|---|---|---|
| $x$ | $\partial_x B = \frac{\partial ln(B)}{\partial ln(x)}$ | $\partial_x \mathcal{S} = \frac{\partial \mathcal{S}}{\partial ln(x)}$ |
| $r$ | 1 | 0 |
| $K$ | $\frac{1}{\kappa - 1}$ | $\frac{-\kappa}{\rho - 1}$ |
| $R_0$ | $\frac{-1}{\kappa - 1}$ | $\frac{\kappa}{\rho - 1}$ |
| $a$ | $\frac{1}{\kappa - 1}$ | $\frac{-\kappa}{\rho - 1}$ |
| $h$ | $\frac{-1}{(\rho - 1)(\kappa - 1)}$ | $\frac{\kappa(1-\rho) + 2\rho}{(\rho - 1)^2}$ |
| $J$ | $\frac{\rho}{(\rho - 1)(\kappa - 1)} + 1$ | $\frac{-2\rho}{(\rho - 1)^2}$ |
| $e$ | $\frac{\rho}{(\rho - 1)(\kappa - 1)} + 1$ | $\frac{-2\rho}{(\rho - 1)^2}$ |

| | | |
|---|---|---|
| $m$ | $\frac{-\rho}{(\rho-1)(\kappa-1)} - 1$ | $\frac{2\rho}{(\rho-1)^2}$ |

Table S3. Sensitivities of biomass ratio and stability to all parameters, including maximum consumption rate, $J$, and half-saturation density, $R_0$.

Thus, we can see that:

$\partial_K B = \partial_a B = -\partial_{R_0} B$ and $\partial_K \mathcal{S} = \partial_a \mathcal{S} = -\partial_{R_0} \mathcal{S}$

$\partial_e B = \partial_J B = -\partial_e B$ and $\partial_e \mathcal{S} = \partial_J \mathcal{S} = -\partial_e \mathcal{S}$

Therefore, our results in the main text directly apply to the Michaelis-Menten formulation of the functional response. We also see that when using $J$ and $R_0$ the sensitivities can be reduced to two groupings, which simplifies analyses further.

**Supplementary Information 4: Stability metric adjustment**

In our study, we qualified stability in terms of the steady state being a stable equilibrium or a limit cycle (oscillations around unstable equilibrium). To this end we used a metric which quantifies the tendency for oscillations in consumer-resource dynamics (Johnson & Amarasekare 2015). The metric was defined as $\phi = ahK$; the smaller the value the more 'stable' the dynamics. In particular, if $\phi < 1$ the dynamics are stable, if $\phi > 1$ they are unstable. Effectively, this metric was deduced as, what its authors referred to, a 'conservative' criterion for stability (for details see derivation of eq. (5) in (Johnson & Amarasekare 2015).

Expressed in terms of the aggregates $\rho$ and $\kappa$, $\phi = \frac{\kappa}{\rho-1}$. We adjusted $\phi = 1$ to create a metric, $\mathcal{S}$, which vanished ($\mathcal{S} = 0$) at the Hopf bifurcation, was positive for the stable equilibrium ($\mathcal{S} > 0$) and negative for oscillations around the unstable equilibrium ($\mathcal{S} < 0$). The Hopf bifurcation in terms of $\phi$ is given by $\phi_H = \frac{\rho+1}{\rho-1}$. Therefore, for $\mathcal{S}$ to vanish at the Hopf bifurcation, we took $\mathcal{S} = \phi_H - \phi = \frac{\rho+1}{\rho-1} - \frac{\kappa}{\rho-1} = \frac{-(\kappa-\rho-1)}{\rho-1}$. Thus, $\mathcal{S} = 0$at the Hopf bifurcation, $\mathcal{S} > 0$ when the equilibrium is stable and $\mathcal{S} < 0$ when the oscillations around the unstable equilibrium occur.

**Supplementary Information 5: Sensitivity analysis methodology**

In the first subsection we clarify our sensitivity analysis methods in relation to the case of correlated parameter changes and the adjustment of the elasticity metric (SI 5.1). In the second subsection, we present the full sensitivity ranking regions in the $\rho$-$\kappa$ plane (SI 5.2).

**5.1: Methodological clarifications on sensitivity analysis**

Two points regarding our sensitivity analysis require additional clarification. First, we show how our method can account for potential correlations among model parameters. Second, we demonstrate why our results remain unaffected by the choice of an adjusted (in place of standard) elasticity as our sensitivity metric for stability.

Assume our response variable, $\mathcal{V}$, is a function of parameters $x_1, x_2, \ldots x_i$. $\mathcal{V}$ applies to any model response variable such as biomass ratio, $\mathcal{B}$, and the stability metric, $\mathcal{S}$. In our approach (main text, '*The dual approach – Sensitivity analysis*'), we quantify how small changes in the model parameters – occurring one parameter at a time - translate into changes in the variable $\mathcal{V}$ (Fig. S5.1, orange arrows). This constitutes a sensitivity analysis. By definition it is expressed in terms of the first-order derivative of $\mathcal{V}$ with respect to a parameter, $x_i$, $\partial\mathcal{V}/\partial x_i$, and it is calculated assuming one parameter is perturbed at a time (Caswell 2019). Standard practices of sensitivity analysis do not include higher-order terms (Chitnis et al. 2008, Manlik et al. 2018, Caswell 2019, Zhao et al. 2020). We should note that a simple sensitivity will use the first derivative directly, while other metrics (e.g., elasticity) may scale the first derivative.

If the parameters $x_1$, $x_2$, ... $x_i$ depend on environmental conditions, $\mathcal{T}$ (e.g. temperature), then changes in $\mathcal{T}$ will induce changes in the variable $\mathcal{V}$. Moreover, the parameter changes with respect to changes in $\mathcal{T}$, $dx_i/d\mathcal{T}$ (Fig. S5.1, green arrows), may be correlated. The sensitivity of $\mathcal{V}$ with respect to temperature, $d\mathcal{V}/d\mathcal{T}$, can be represented as a linear combination of sensitivities of $\mathcal{V}$ with respect to the individual parameters, $\partial\mathcal{V}/\partial x_i$ (Fig S5.1, orange arrows). The coefficients of this linear combination are the derivatives of the parameters with respect to temperature, $dx_i/d\mathcal{T}$ (Fig S5.1, green arrows). Importantly, the linear combination accounts for the potential correlations between changes in parameter values induced by a change of temperature. We present this argument in the equations below:

We have that

$$\mathcal{V} = \mathcal{V}(x_1(\mathcal{T}), x_2(\mathcal{T}), ...) \quad \text{(S5.1)}$$

The sensitivity of $\mathcal{V}$ with respect to $\mathcal{T}$ is defined as

$$d\mathcal{V}/d\mathcal{T} = d\mathcal{V}(x_1(\mathcal{T}), x_2(\mathcal{T}), ...) /d\mathcal{T} \quad \text{(S5.2)}$$

By applying the chain rule this becomes

$$d\mathcal{V}/d\mathcal{T} = \partial\mathcal{V}/\partial x_1 * dx_1/d\mathcal{T} + \partial\mathcal{V}/\partial x_2 * dx_2/d\mathcal{T} + ... + \partial\mathcal{V}/\partial x_i * dx_i/d\mathcal{T} \quad \text{(S5.3)}$$

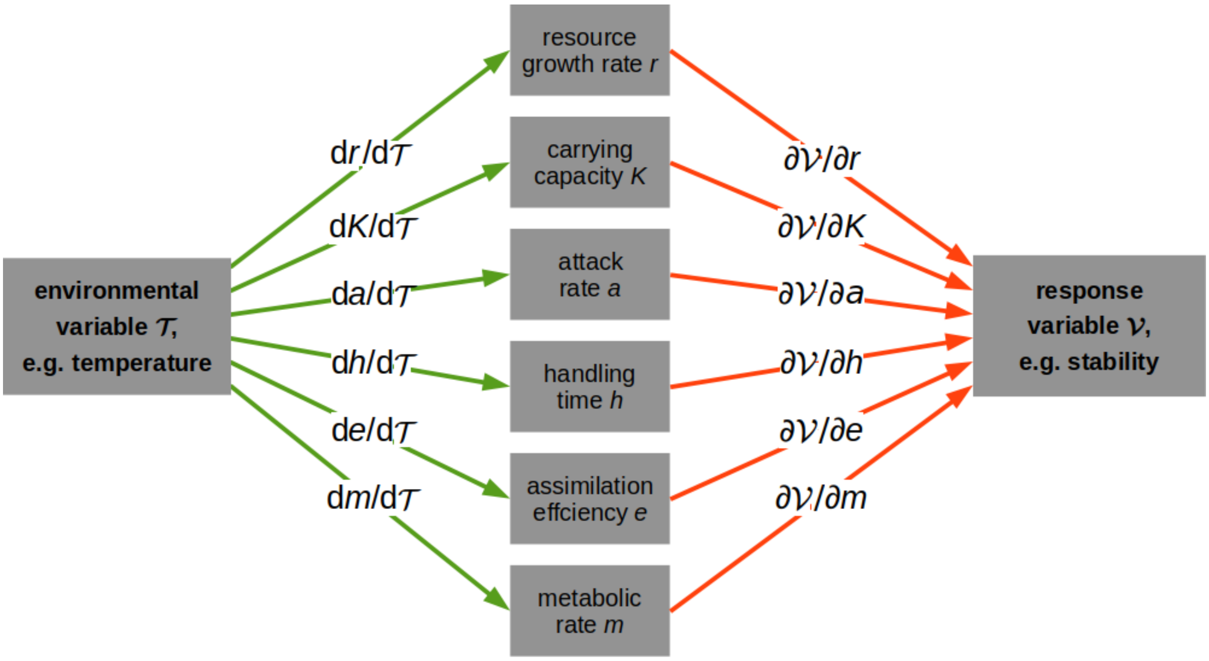


Figure S5.1. A graphical representation of equation S5.3. The sensitivity of the model variable, $\mathcal{V}$, with respect to changes in the individual parameters, $\partial\mathcal{V}/\partial x_i$ (orange arrows) quantifies each parameter's isolated impact on $\mathcal{V}$, which can be measured with different sensitivity metrics (e.g., elasticity). If the parameter values (and ultimately the variable $\mathcal{V}$) depend on some environmental condition, $\mathcal{T}$ (e.g., temperature), then changes in temperature will induce simultaneous changes in these parameters. These changes are quantified by the first derivative of each parameter with respect to temperature, $dx_i/d\mathcal{T}$ (green arrows). Thus, the impact of small temperature changes on the variable $\mathcal{V}$, $d\mathcal{V}/d\mathcal{T}$, is determined by a weighted sum of the individual effects of all parameters, i.e., the sum of the combined green (weight, $dx_i/d\mathcal{T}$) and orange (sensitivity, $\partial\mathcal{V}/\partial x_i$) arrows.

This formula can be written in terms of elasticities as well:

$$d\ln(\mathcal{V})/d\ln(\mathcal{T}) = \mathcal{T}/\mathcal{V} * \partial\mathcal{V}/\partial\mathcal{T}$$

$$= \mathcal{T} * dx_1/d\mathcal{T} * 1/\mathcal{V} * \partial\mathcal{V}/\partial x_1 + \mathcal{T} * dx_2/d\mathcal{T} * 1/\mathcal{V} * \partial\mathcal{V}/\partial x_2 + \ldots$$

$$= \mathcal{T}/x_1 * dx_1/d\mathcal{T} * x_1/\mathcal{V} * \partial\mathcal{V}/\partial x_1 + \mathcal{T}/x_2 * dx_2/d\mathcal{T} * x_2/\mathcal{V} * \partial\mathcal{V}/\partial x_2 + \ldots$$

$= \mathrm{dln}(x_1)/\mathrm{dln}(\mathcal{T}) * \partial\ln(\mathcal{V})/\partial\ln(x_1) + \mathrm{dln}(x_2)/\mathrm{dln}(\mathcal{T}) * \partial\ln(\mathcal{V})/\partial\ln(x_2) + \ldots$

$= \mathrm{dln}(x_1)/\mathrm{dln}(\mathcal{T}) * \partial_{x1}\mathcal{V} + \mathrm{dln}(x_2)/\mathrm{dln}(\mathcal{T}) * \partial_{x2}\mathcal{V} + \ldots$

where $\partial_{xi}\mathcal{V}$ is the elasticity of $\mathcal{V}$ with respect to $x_i$ and $\mathrm{dln}(x_i)/\mathrm{dln}(\mathcal{T})$ measures the dependence of $x_i$ on $\mathcal{T}$ on a double logarithmic scale. We conclude that the one-parameter elasticities studied in the main text are the building blocks to study the elasticities for correlated parameter changes.

With respect to the adjusted sensitivity metric used in the stability sensitivity analysis, we argue here why its application does not alter the results. We used two variants of elasticity as our sensitivity metric. For the biomass ratio, $B$, we used the standard elasticity, $\partial_x B = \frac{\frac{\partial B}{B}}{\frac{\partial x}{x}} = \frac{\partial ln(B)}{\partial ln(x)}$ (Fig. S5.2). While for stability, $\mathcal{S}$, we adjusted this to not include the relative change in $\mathcal{S}$, i.e., $\partial_x \mathcal{S} = \frac{\partial \mathcal{S}}{\frac{\partial x}{x}} = \frac{\partial \mathcal{S}}{\partial ln(x)}$ (Fig. S5.3). There exist two reasons for this adjustment. First, the Hopf bifurcation occurs at $\mathcal{S} = 0$. As $\mathcal{S} \rightarrow 0$, elasticities diverge, $\frac{\partial ln(\mathcal{S})}{\partial ln(x)} \rightarrow \infty$ (Fig. S5.4). Our adjusted metric does not have these divergences but preserves the main properties of the standard elasticity metric. In particular, the rankings of stability sensitivity across the $\rho - \kappa$ plane (Figs 5a-b, 6a-c, and 7a-f in the main text) are identical for the standard and alternative elasticity metric. The reason for this equivalence is that the standard elasticity metric is retrieved by dividing our adjusted metric by stability, $\mathcal{S}$. This division only scales the elasticity metric (the same scaling for all parameters), and does not change the ranking, or any of our conclusions. Additionally, the comparison of the dependence of the two stability metrics on the aggregate parameters (and hence on the individuals parameters; Figs S5.3 and S5.4) shows that our adjusted elasticity metric has a much smoother dependence than the standard one (due to the divergences of the latter).

Second, stability switches from stable ($\mathcal{S} < 0$) to unstable ($\mathcal{S} > 0$) at the Hopf bifurcation. In case of using elasticity, which only quantifies the relative change in $\mathcal{S}$, if a parameter was destabilising (e.g. attack rate) it would have positive sensitivity in the region where $\mathcal{S} < 0$ ($\mathcal{S}$ would become more negative, i.e., destabilising effect) and a negative sensitivity when $\mathcal{S} > 0$ ($\mathcal{S}$ would decrease, i.e., destabilising) (Fig. S5.4). This, combined with the divergence around the Hopf bifurcation, render the analysis based on the elasticity of stability very hard to follow. Through our adjustment we avoid both issues: we do not divide by 0, and by looking at the absolute change in $\mathcal{S}$ caused by a relative parameter perturbation, we always get an indication of whether $\mathcal{S}$ increases or decreases. Most significantly, this adjustment does not alter the sensitivity relationships, or the patterns observed.

Below we present an illustration of how we derive the sensitivity formulas in Table 1 of the main text. Following this, we plot $\partial_x B$ (Fig. S5.1), $\partial_x \mathcal{S}$ (Fig. S5.2) and the unadjusted stability elasticity (Fig. S5.3) with respect to all six parameters (i.e., contour plots of the values in Table 1 in the $\rho - \kappa$ plane).

Illustration of derivation of biomass elasticity with respect to attack rate, $a$:

$$\partial_a B = \frac{\frac{\partial B}{B}}{\frac{\partial x}{a}} = \frac{\partial B}{\partial}\frac{a}{B} = \frac{er}{a^2K(e-mh)}\frac{ma^2K(e-mh)}{er(aeK-ahKm-m)} = \frac{m}{m\left(ahK\left(\frac{e}{mh}-1\right)-1\right)} = \frac{1}{\kappa-1}$$

Similarly, for stability sensitivity with respect to attack rate, $a$:

$$\partial_a \mathcal{S} = \frac{\partial \mathcal{S}}{\frac{\partial a}{a}} = \frac{\partial \mathcal{S}}{\partial a} a = -hKa = \frac{-\kappa}{\rho - 1}$$

Below are the contour plots of the sensitivities for biomass ratio (Fig. S5.2) and stability (Fig. S5.3). Fig. S5.4 shows the standard elasticity of the stability metric.

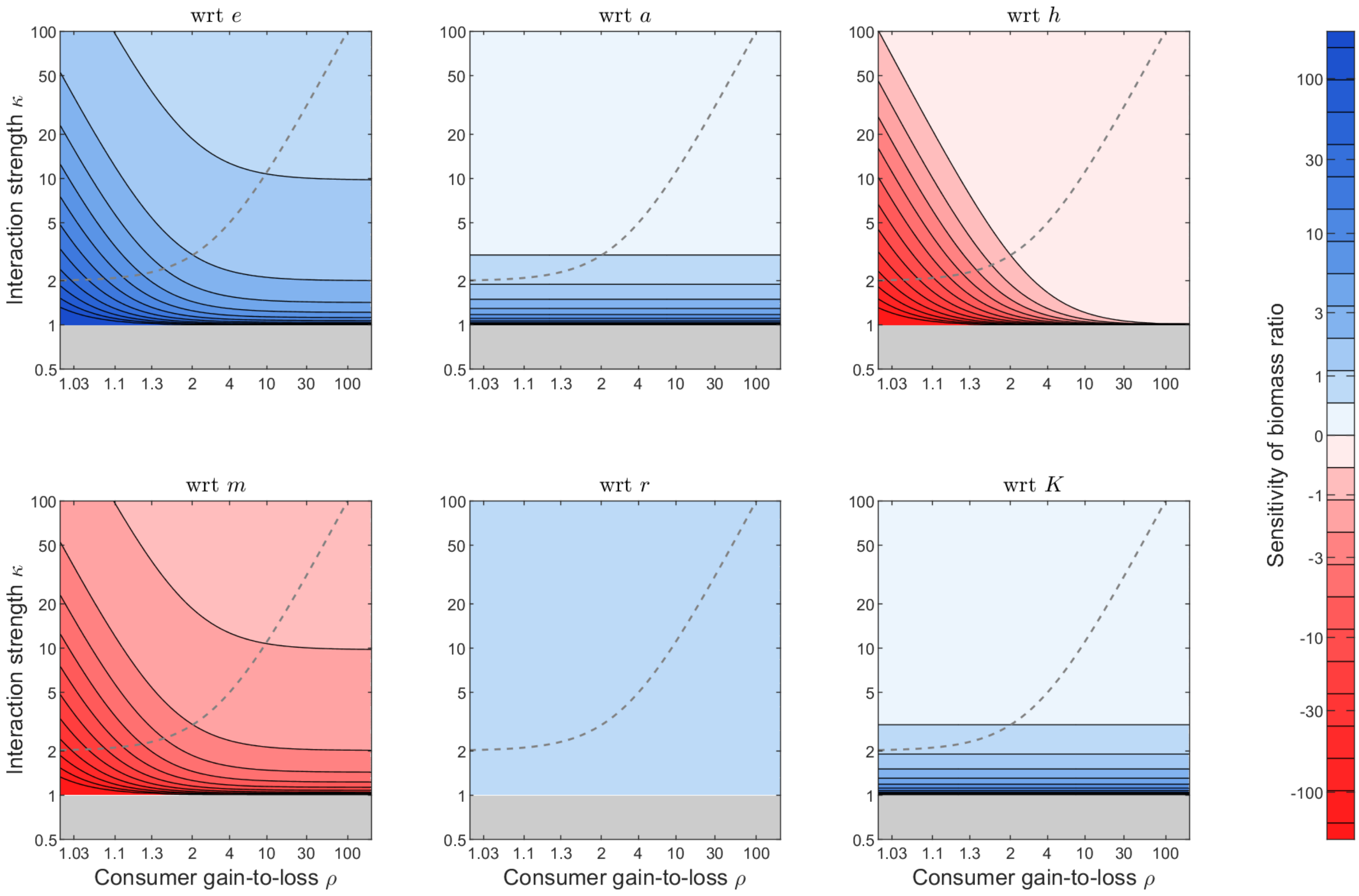


Figure S5.2. Values of biomass ratio sensitivities with respect to ('wrt') individual parameters (different panels) derived from the expressions in Table 1 of the main manuscript.

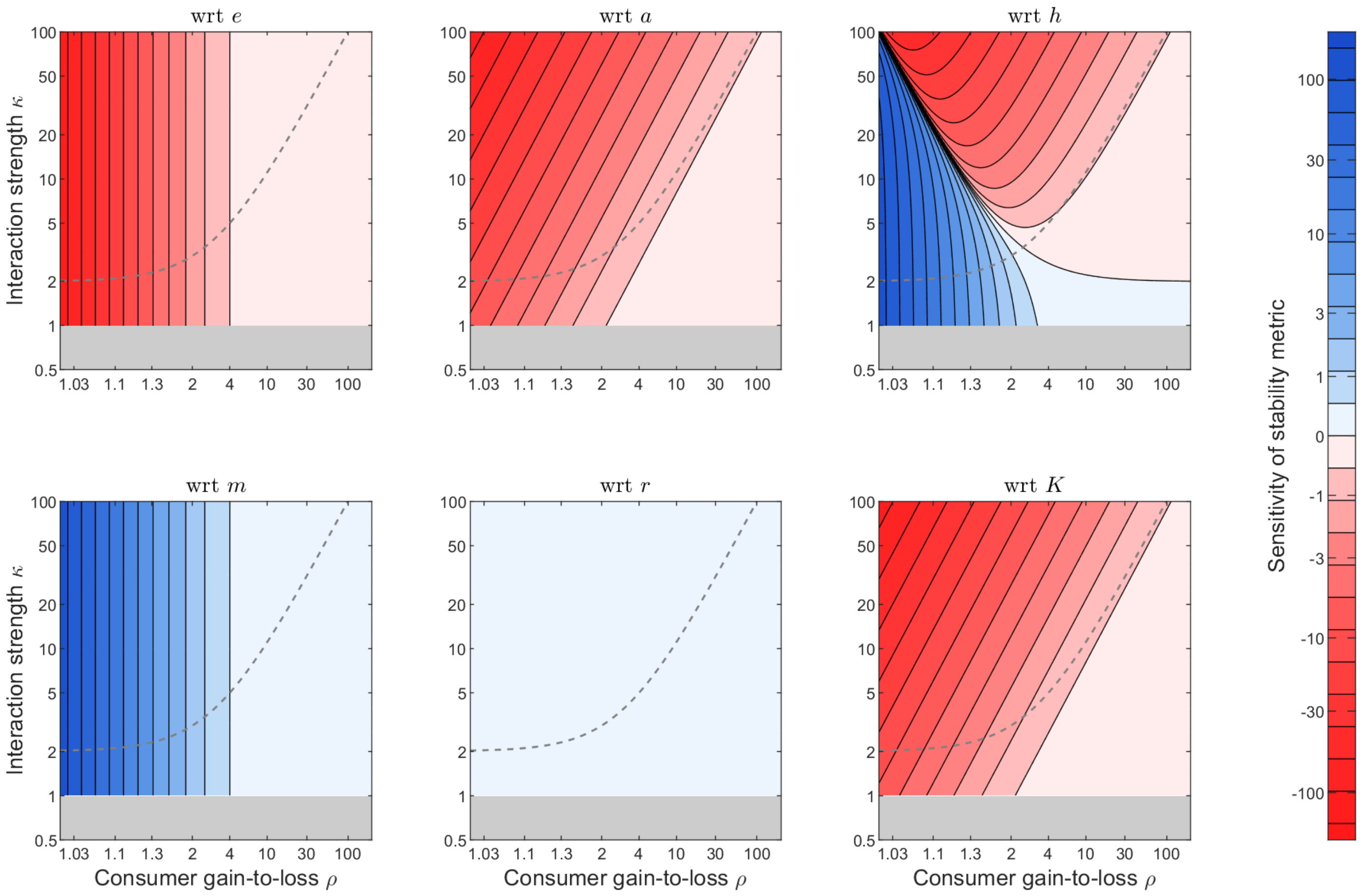


Figure S5.3. Values of stability metric sensitivities with respect to (‘wrt’) to individual parameters (different panels) derived from the expressions in Table 1 of the main manuscript.

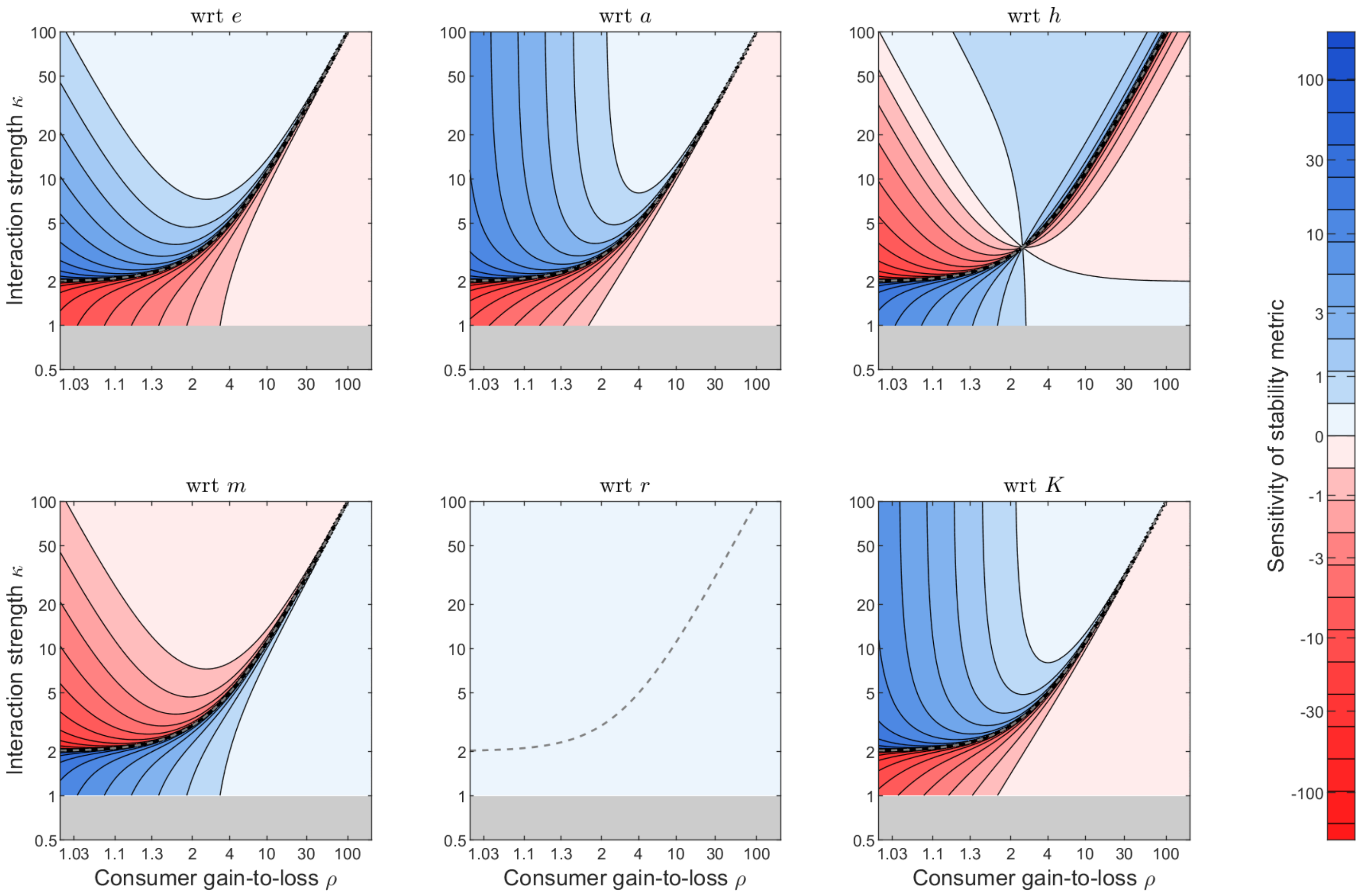


Figure S5.4. Values of the sensitive metric elasticities, i.e., the unadjusted sensitivity metric with respect to ('wrt') to individual parameters (different panels). Elasticity (i.e., relative change in $\mathcal{S}$) contours become much harder to read compared to the adjusted elasticity (Fig. SI5.3, absolute change in $\mathcal{S}$) for two reasons: i) approaching the Hopf bifurcation (diagonal dashed line) all elasticities diverge as we divide by 0, distorting the contours, ii) the sign of the contours switches as they cross the Hopf bifurcation, even though the effect of the parameters on stability does not change.

### 5.2 The full sensitivity ranking regions in the *ρ*-*κ* plane

The sensitivity expressions in Table 1 of the main text can be ordered in terms of their magnitude. These orderings create conditions which can be plotted onto the $\rho$-$\kappa$ plane and split it in different regions. In each region a unique ranking of all sensitivities occurs. For the biomass ratio (Fig. S5.5) there are six regions. This occurs because $\partial_e B = -\partial_m B$ always rank first and $\partial_a B = \partial_K B$. So, there were three elasticities to be ranked: $\partial_a B$, $\partial_r B$ and $\partial_h B$, creating six possible combinations of their rankings. The stability metric had four uniquely-defined regions (Fig. S5.6). In this case $-\partial_e B = \partial_m B$, $\partial_a B = \partial_K B$ and $\partial_r B = 0$, reduced the number of sensitivities to be ranked down to three. However, $\partial_h B$ never becomes the largest sensitivity, and hence either $|\partial_e B \vee$ or $\partial_a B \vee$ can rank first. This leaves two alternative orderings per scenario, producing a total of four regions.

12

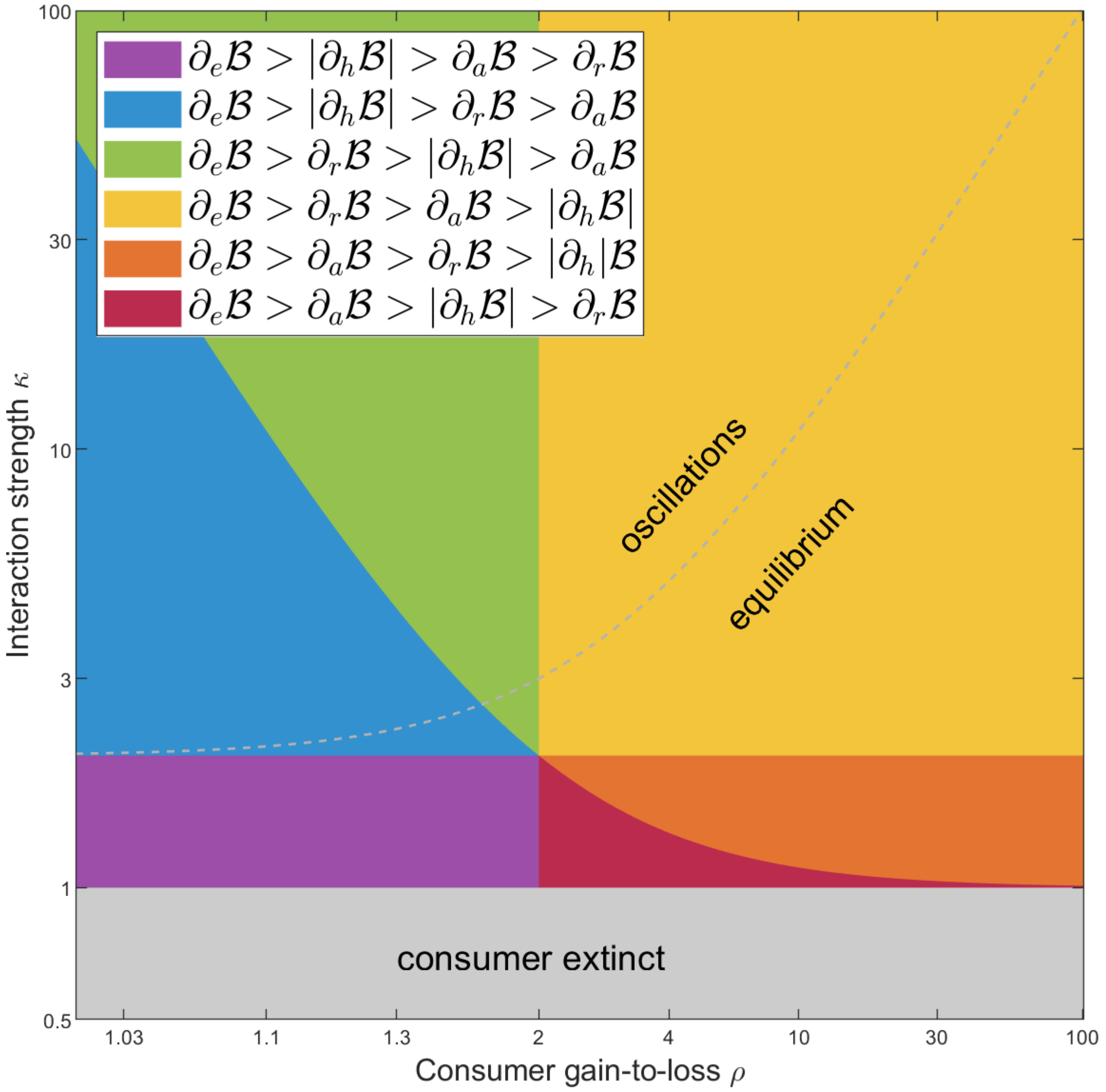

Figure S5.5. Biomass ratio elasticities ranked in the $\rho$-$\kappa$ plane. The six regions correspond to unique rankings of the elasticities. Close to consumer extinction ($\kappa$=1) either $|\partial_h B|$ (purple region) or $\partial_a B = \partial_K B$ (red region) rank second. Moving away from the consumer boundary, $\partial_r B$ increases and becomes the second highest elasticity (yellow and green regions). In Fig. 3a from the main text, the regions where the top-two ranked elasticities remain unchanged (purple and blue, green and yellow, orange and red) have been merged into single regions, ignoring changes in the lowest-ranking elasticities.

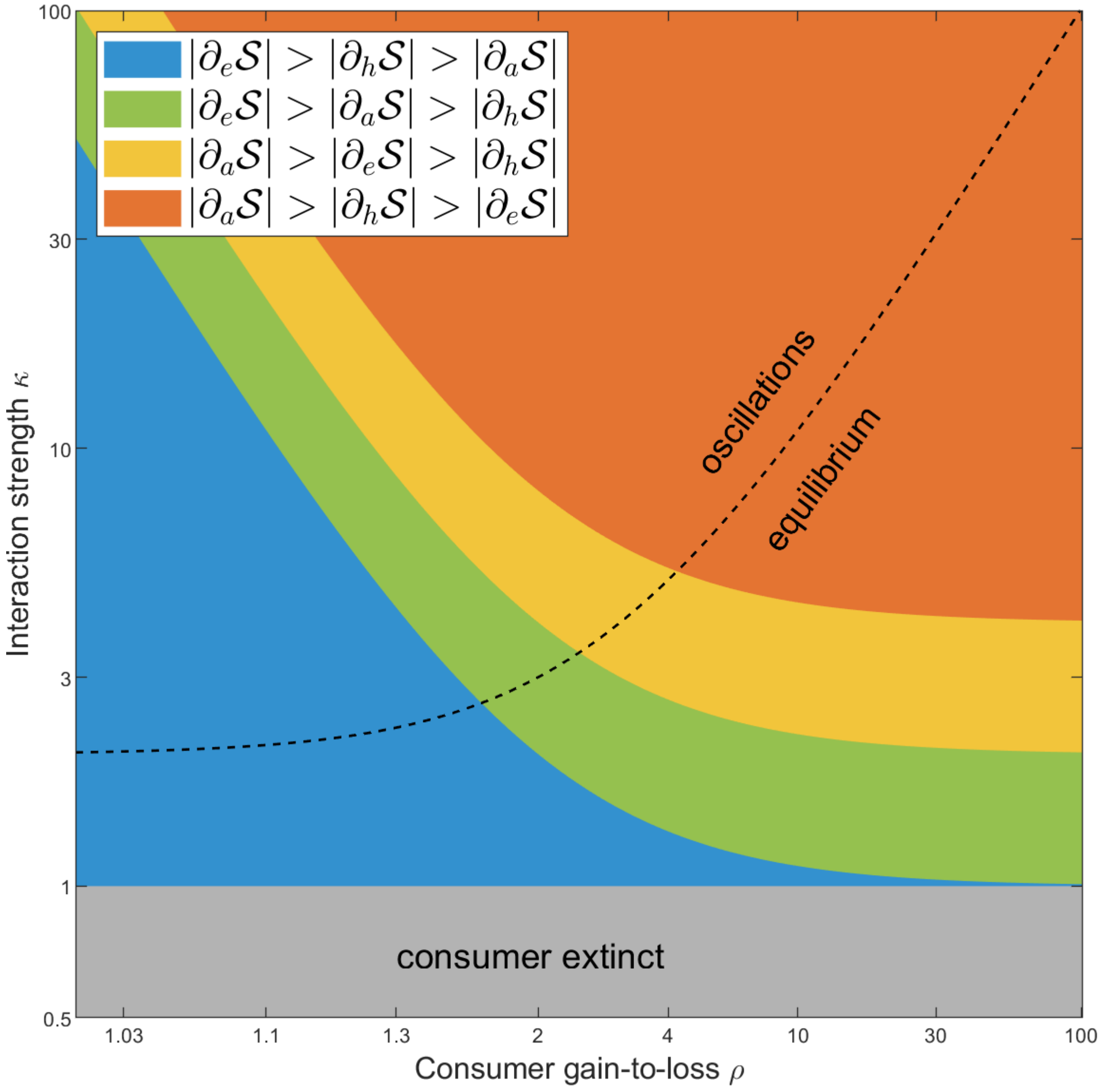

Figure S5.6. Stability metric sensitivities ranked in the $\rho$-$\kappa$ plane. The four regions correspond to unique rankings of the elasticities. Close to consumer extinction ($\kappa$=1) $|\partial_e \mathcal{S}| = \partial_m \mathcal{S}$ (blue region) rank first, $|\partial_h \mathcal{S}|$ second and $|\partial_a \mathcal{S}| = |\partial_K \mathcal{S}|$ third. Moving away from the consumer boundary, $|\partial_a \mathcal{S}| = |\partial_K \mathcal{S}|$ increases, first switching rankings with $|\partial_h \mathcal{S}|$ (green region) and then with $|\partial_e \mathcal{S}| = \partial_m \mathcal{S}$ to become the top sensitivity (yellow region). Furthest from consumer extinction, $|\partial_h \mathcal{S}|$ becomes the second highest sensitivity (orange region).

**Supplementary Information 6: Temperature parameterisations from the literature**

In our study, we used the Rosenzweig-MacArthur model (Rosenzweig & MacArthur 1963) to derive the framework combining a sensitivity analysis and the aggregate parameters, maximal energetic efficiency, $\rho$, and interaction strength, $\kappa$. We argued that the framework can be applied to any temperature parameterisation which can be fitted to the Rosenzweig-MacArthur model. We implemented parameterisations from six studies from the literature with empirically-derived thermal dependencies of certain parameters and assumptions about the thermal dependence of others. We should note that not all studies measured or reported a thermal performance curve for the resource growth rate, $r$. Therefore, the consumer-resource biomass ratio (eqs. 3 and S1.5) was evaluated and presented only for two studies (Fig. 2). However, biomass ratio sensitivities can be expressed in terms of $\rho = \frac{e}{mh}$ and $\kappa = ahK\left(\frac{e}{mh} - 1\right)$ and hence no$r$ thermal dependence is necessary to plot the parameterisation trajectories in the $\rho - \kappa$ plane. As such, all six studies utilised in our application of the framework (Fig. 4) reported parameterisations enabling us to derive the thermal dependencies of $\rho$ and $\kappa$. Finally, we note that these parametrisations all assumed a type II functional response. Below we briefly describe each study before providing details of how we implemented their parameterisations.

*Brief description of studies:*

Uszko *et al.* (2017)investigated the effects of warming on the stability of consumer-resource communities using the Rosenzweig-MacArthur model. Their model was parameterised for the crustacean *Daphnia hyaline* feeding on the green alga *Monoraphidium minutum*.

Fussmann *et al.* (2014) combined a meta-analysis of global invertebrate data and literature sources to determine the thermal dependencies of biological rates implemented in a bioenergetic model (Yodzis & Innes 1992b). The functional response was represented in terms of maximum consumption rate, $J$, and resource half-saturation density, $R_0$ (see Supplementary Information 3).

Vucic-Pestic *et al.* (2011): an empirical study of how the functional response (attack rate, handling time) and consumer metabolism scale with temperature in a predator-prey community with three different size-classes of predators and two types of prey (mobile and resident). Other parameter values were taken from the literature. No model was employed, though the energetic efficiency term was based on Vasseur and McCann (2005).

Binzer *et al.* (2016) performed a modelling study of the interactive effects of eutrophication and warming on food webs parameterised from the literature. The population dynamics between successive trophic levels were described using a bio-energetic model (Yodzis & Innes 1992b).

Sentis *et al.* (2012): experimental data from a ladybeetle (*C. Maculata lengi*) feeding on a green peach aphid (*Myzus Persicae*) used to estimate how the functional response scaled with temperature. No model was employed, though the energetic efficiency term was based on Vasseur and McCann (2005).

Archer *et al.* (2019): a comparison of the thermal dependence of the functional responses derived from field and lab measurements. A sedentary (larvae of *L. riparia*) and a mobile (larvae of *P. cingulatus*) predator fed on a common prey (blackfly larvae from the *Simuliidae* family). A per capita energy feeding rate was derived, which was then transformed to the dimensionless energetic efficiency – as in the previous studies.

Parameter values in studies:

In the following equations, temperature $T$ is in Kelvin, where 273.15K corresponds to 0°C. In all Arrhenius-based parameterisations, $k = 8.62 \times 10^{-5}$ $[eVK^{-1}]$ is the Boltzmann constant. $r$is the resource population growth rate (time$^{-1}$), $a$ is the attack or search rate (surface or volume per unit of time and resource unit), $h$ is the handling time (time), $m$ is the metabolic rate (time$^{-1}$) or $m_E$ the standard metabolic rate (energy*time$^{-1}$), $J$ is the maximum feeding rate (time$^{-1}$), $R_0$ the resource-half saturation density (biomass or resource unit per surface or volume), $K$ is the resource carrying capacity (biomass or resource unit per surface or volume) and $e$ is the consumer assimilation efficiency (unitless).

<u>Uszko et al. (2017):</u>

$r(T) = r_0 e^{\frac{-(T-T_{opt})^2}{2s^2}}$ with $r_0 = 2.2, T_{opt} = 298.15, s = 12$, $[d^{-1}]$

$a(T) = a_0 e^{\frac{-(T-T_{opt})^2}{2s^2}}$ with $a_0 = 6, T_{opt} = 296, s = 9.4$, $[m^3 g^{-1} d^{-1}]$

$h(T) = h_0 e^{\frac{(T-T_{opt})^2}{2s^2}}$ with $h_0 = 0.2, T_{opt} = 294.1, s = 7.2$, $[d]$

$m(T) = m_0 e^{\frac{-E_m}{kT}}$with $m_0 = 4.4 * 10^8, E_m = 0.55$, $[d^{-1}]$

$K$ and $e$ are not functions of temperature; we used $e = 0.385$ and $K = 2$, $[gm^{-3}]$.

Fussmann et al. (2014)

Reference temperature $T_{ref} = 20°C = 293.15K$. Carnivore assimilation efficiency was used: $e = 0.85$

$r(T) = r_0 e^{\frac{E_r(T-T_{ref})}{kTT_{ref}}}$ with $r_0 = 8.715 * 10^{-7}, E_r = 0.84$, $[s^{-1}]$

$K(T) = K_0 e^{\frac{E_K(T-T_{ref})}{kTT_{ref}}}$ with $K_0 = 5.623, E_K = -0.772$, $[gm^{-2}]$

$R_0(T) = R_{0_0} e^{\frac{E_{R_0}(T-T_{ref})}{kTT_{ref}}}$ with $R_{0_0} = 3.664, E_{R_0} = -0.114$, $[gm^{-2}]$

$J(T) = J_0 e^{\frac{E_J(T-T_{ref})}{kTT_{ref}}}$ with $J_0 = 8.40810^{-6}, E_J = 0.467$, $[s^{-1}]$

$m(T) = m_0 e^{\frac{E_m(T-T_{ref})}{kTT_{ref}}}$ with $m_0 = 2.689 * 10^{-6}, E_m = 0.639$, $[s^{-1}]$

Vucic-Pestic et al. (2011):

$T_{ref} = 290.65K$ and $e = 0.85$.

Predators were split into three mass classes.Predator masses within the same size-classes varied with temperature, so we used the mean value over all temperatures to obtain the mean mass of each size class. Additionally, predator masses varied between the experiments for the two prey types.

| Prey mass ($m_R$) $[mg]$ | 1.91 (mobile) | 23.26 (resident) |
| --- | --- | --- |

| | | |
|---|---|---|
| Small predator mass ($m_C$) [$mg$] | 66.1617 | 72.6217 |
| Medium predator mass ($m_C$) [$mg$] | 121.2167 | 134.8133 |
| Large predator mass ($m_C$) [$mg$] | 147.8683 | 160.6550 |

Parameter thermal dependencies also scale with body mass. For the dimensions in the equations to be correct, we introduce a reference body mass for consumers and resources, $m_{C_0} = 1mg$ and $m_{R_0} = 1mg$, respectively.

We also note that the study does not explicitly measure or define a resource carrying capacity. However, it does provide an equation of the thermal dependence of the prey population. We assumed this to be the carrying capacity in our calculation of $\kappa$.

Finally, in this study, the authors measure the standard metabolic rate with units of energy over time [$Jh^{-1}$]. For the calculation of maximal energetic efficiency, the maximal feeding rate [$\frac{1}{handlingtime}$] was converted from [$\frac{1}{time}$] to an energetic equivalent, [$Jh^{-1}$]. The conversion – described in the original study - is from per capita to per capita biomass (multiply with prey biomass $m_R$) and then to energy per capita (multiply with the weight [$kg$] - energy [$J$] conversion factor of $7 \times 10^6$). Note,$m_R = [mg] = 10^{-6}$].

$$\rho = 7 \times 10^6 \times 10^{-6} \times m_R \times \frac{1}{h} \times \frac{e}{m}$$

Similarly, for $\kappa = ahK\left(\frac{e}{mh} - 1\right)$, the first term required a correction, since $h$ needed a conversion from per capita $[hInd^{-1}]$ to per capita biomass time $[hmg^{-1}]$ to match the dimensions of $K$. Hence, we divided handling time by the prey biomass:

$$\kappa = a\left(\frac{1}{m_R}h\right)K(\rho - 1)$$

The parameter values we used for calculating $\rho$ and$\kappa$:

The carrying capacity:

$$K(T) = K_0\left(\frac{m_R}{m_{R_0}}\right)^s e^{\frac{-E_K}{kT}}\sigma^z e^{tl_0(tl-1)}$$

where $K_0 = e^{-31.15}$, $m_R$ is the prey body mass,$s = -0.72$, $E_K = -0.71$, $z = 1.03$, $tl_0 = 2.86$, $tl = 1.5$.

$\sigma = \sigma_0 e^{\frac{-E_\sigma(T_{ref}-T)}{kTT_{ref}}}$ $[gCm^{-2}y^{-1}]$, where $\sigma_0 = 600$ and $E_\sigma = 0.35$.

Attack rate $[m^2h^{-1}]$ and handling time $[hInd^{-1}]$ varied with prey type:

$$a(T) = a_0\left(\frac{m_C}{m_{C_0}}\right)^s e^{\frac{-E_a(T_{ref}-T)}{kTT_{ref}}}$$

with:

$a_0 = e^{-1.77}, E_a = 0.37, s = -0.48$ for mobile prey

$a_0 = e^{-5.94}, E_a = -0.27, s = 0.91$ for resident prey

$$h(T) = h_0 \left(\frac{m_C}{m_{C_0}}\right)^s e^{\frac{-E_h(T_{ref}-T)}{kTT_{ref}}}$$

with:

$h_0 = e^{2.99}, E_h = -0.24, s = -0.66$ for mobile prey and

$h_0 = e^{6.85}, E_h = -0.23, s = -1$ for resident prey

Metabolic rate was measured in terms of energy per unit of time, $m_E(T)$.

$m_E(T) = m_0 \left(\frac{m_C}{m_{C_0}}\right)^s e^{\frac{-E_m(T_{ref}-T)}{kTT_{ref}}}$ with $m_0 = e^{-3.91}, E_m = 0.61, s = 0.62$, with units $[Jh^{-1}]$.

$m_C$ varieswith the predator size-class.

Binzer et al. (2016):

$m_R$ and $m_C$ are the unit biomass $[g]$ of prey and predator, respectively. For the dimensions in the following equations to be correct, we introduce a reference body mass for resource and consumer, $m_{R_0} = 1g$ and $m_{C_0} = 1g$, respectively. We assumed a 100:1 consumer-resource body-mass ratio. $T_{ref} = 20°C = 293.15K$ and $e = 0.85$.

$r(T) = r_0 \left(\frac{m_R}{m_{R_0}}\right)^s e^{\frac{E_r(T_{ref}-T)}{kTT_{ref}}}$ with $r_0 = e^{15.68}, E_r = -0.84, s = -0.25$, $[s^{-1}]$

$K(T) = K_0 e^{\frac{E_K(T_{ref}-T)}{kTT_{ref}}}$ with $E_K = 0.71$, $[gm^{-2}]$

In the original study $K_0$ varied between 1 and 20, so we selected two scenarios, $K_0 = 5$ for ‘low enrichment’ and $K_0 = 15$ for ‘high enrichment’ (Fig. 4c, 7c).

$$a(T) = a_0 \left(\frac{m_R}{m_{R_0}}\right)^{s_R} \left(\frac{m_C}{m_{C_0}}\right)^{s_C} e^{\frac{E_a(T_{ref}-T)}{kTT_{ref}}} \text{with } a_0 = e^{13.1}, E_a = -0.38, s_R = 0.25, s_C = -0.8,$$

$[m^2 s^{-1}]$

$$h(T) = h_0 \left(\frac{m_R}{m_{R_0}}\right)^{s_R} \left(\frac{m_C}{m_{C_0}}\right)^{s_C} e^{\frac{E_h(T_{ref}-T)}{kTT_{ref}}} \text{with } h_0 = e^{9.66}, E_h = 0.26, s_R = -0.45, s_C = 0.47, [s]$$

$$m(T) = m_0 \left(\frac{m_C}{m_{C_0}}\right)^{s} e^{\frac{E_m(T_{ref}-T)}{kTT_{ref}}} \text{with } m_0 = e^{16.54}, E_m = -0.69, s = -0.31, [s^{-1}]$$

Sentis et al (2012):

Certain parameterisations were not fully provided in the original manuscript. We have retrieved them from the authors and used these for the calculation of maximal energetic efficiency, $\rho$, and interaction strength, $\kappa$.

The prey and predator masses are $m_R = 0.5031\ [mg]$ and $m_C = 8.7 \times 10^{-3} [g]$, respectively. For the dimensions in the parameter equations to be correct, we introduce a reference body mass for the consumer, $m_{C_0} = 1g$.

We note that this study too does not explicitly measure or define a resource carrying capacity. However, it does provide a gradient of prey densities based on which their energetic efficiency was calculated. We used a low and a high value from the gradient as our carrying capacity values.

Similarly to Vucic-Pestic *et al.* (2011), standard metabolic rate was measured [energy*time$^{-1}$]. Hence we needed to convert $h$ from $[dInd^{-1}]$ into an energetic equivalent for its units to cancel out with $m$ $[Js^{-1}]$. We applied the conversion described in the original study, to per capita biomass (multiply with $m_R$) and then to energy per capita (multiply with the weight $[kg]$ - energy $[J]$ conversion factor of $7 \times 10^6$). Note,$m_R = [mg] = 10^{-6}]$. Additionally, we scaled $m$ up from seconds to a daily rate.

$$\rho = 7 \times 10^6 \times 10^{-6} \times m_R \times \frac{1}{h} \times \frac{1}{86400m} \times e$$

$\kappa = ahK(\rho - 1)$ did not require any scalings, since the product $ahK$ was unitless.

The parameter values for calculating $\rho$ and $\kappa$ were:

$a(T) = a_0(T - T_0)(T_1 - T)^{\frac{1}{2}}$with $a_0 = 0.061, T_0 = 11.05 + 273.15, T_1 = 38.00 + 273.15,$ $[m^2d^{-1}]$

$h(T) = h_0 e^{\frac{E_h}{T-273.15}}$ with $h_0 = 0.0083, E_h = 44.7569,$ $[dind^{-1}]$

$m_E(T) = m_0 \left(\frac{m_C}{m_{C_0}}\right)^s e^{\frac{-E_m}{kT}}$with $m_0 = 2.86 * 10^7, E_m = 0.65, s = 0.75,$ $[Js^{-1}]$

$K$ $[indm^{-2}]$ corresponnds to prey density values of the original study. We used $K = 5$ and $K = 60$ for the ‘low enrichment’ and ‘high enrichment’ scenarios, respectively (Fig. 4d, 7d).

<u>Archer et al. (2019):</u>

$T_{ref} = 283.15K$

For the dimensions in the parameter equations to be correct, we introduce a reference body mass for the consumer, $m_{C_0} = 1mg$.

No carrying capacity values were explicitly defined or provided. We derived the carrying capacity, $K$, from Fig. 3c of the manuscript which demonstrates the temperature-depedence of the prey density.

For the metabolic rate of *P. cingulatus* we did not include additional the quadratic term used in the fitting of the curve to the original data.

To attain a unitless aggregate parameter $\rho = \frac{e}{mh}$ we converted $h$ from $[dInd^{-1}]$ into an energetic equivalent for its units to cancel out with $m$ $[Jh^{-1}]$ – as in the original study. The per capita handling time was converted to per capita energy handling time using the prey body mass ($0.546mg$) and the prey energy content ($23.1Jmg^{-1}$) - these scaling were provided in the original study. Finally, we converted $m$ from an hourly to a daily rate to match the feeding rate temporal scale.

$$\rho = 0.546 \times 23.1 \times \frac{1}{h} \times \frac{1}{24m} \times e$$

$\kappa = ahK(\rho - 1)$ did not require any scalings, since the product $ahK$ was unitless.

The parameter values used were the following:

For the sedentary (*L. riparia*) predator:

$a(T) = a_0 e^{\frac{-E_a(T_{ref}-T)}{kTT_{ref}}}$ with $a_0 = 0.241$ (Lab 2013), $a_0 = 0.802$ (Lab 2015) or $a_0 = 1.889$ (Field 2015) and $E_a = 0.74$(for all measurements), with units $[m^2 d^{-1}]$

$h = 4.033$, $[d \in d^{-1}]$ – no temperature dependence

$m_E(T) = m_0 \left(\frac{m_C}{m_{C_0}}\right)^s e^{\frac{-E_m}{kT}}$with $m_0 = e^{-4.171} * 10^7, E_m = 0.687, m_C = 10^{0.25}, s = 0.525$, $[Jh^{-1}]$

For the mobile predator (*P. cingulatus*):

$a(T) = a_0 e^{\frac{-E_a(T_{ref}-T)}{kTT_{ref}}}$ with $a_0 = 1.529$ (Lab 2015) or $a_0 = 5.515$ (Field 2015) and $E_a =$ 0.229(for all measurements), with units $[m^2 d^{-1}]$

$h = 0.644$, $[d \in d^{-1}]$ – no temperature dependence

$m_E(T) = m_0 e^{\frac{-E_m}{kT}}$with $m_0 = e^{-1.056} * 10^7, E_m = 1.072$, with no effects of body mass, $[Jh^{-1}]$

And for all calculations:

$K(T) = 10^{0.16(T-273.15)}$, with units $[indm^{-2}]$

$$e(T) = \frac{e_0 e^{\frac{-E_e(T_0 - T)}{kTT_0}}}{e_0 e^{\frac{-E_e(T_0 - T)}{kTT_0}} + 1}, \text{ with } e_0 = e^{2.266}, E_a = 0.164, T_0 = 293.15K$$
